# Which mouse multiparental population is right for your study? The Collaborative Cross inbred strains, their F1 hybrids, or the Diversity Outbred population

**DOI:** 10.1101/2022.08.26.505416

**Authors:** Gregory R. Keele

## Abstract

Multiparental populations (MPPs) encompass greater genetic diversity than traditional experimental crosses of two inbred strains, enabling broader surveys of genetic variation underlying complex traits. Two such mouse MPPs are the Collaborative Cross (CC) inbred panel and the Diversity Outbred (DO) population, which are descended from the same eight inbred strains. Additionally, the F1 intercrosses of CC strains (CC-RIX) have been used and enable study designs with replicate outbred mice. Genetic analyses commonly used by researchers to investigate complex traits in these populations include characterizing how heritable a trait is, *i*.*e*., its heritability, and mapping its underlying genetic loci, *i*.*e*., its quantitative trait loci (QTLs). Here we evaluate the relative merits of these populations for these tasks through simulation, as well as provide recommendations for performing the quantitative genetic analyses. We find that sample populations that include replicate animals, as possible with the CC and CC-RIX, provide more efficient and precise estimates of heritability. We report QTL mapping power curves for the CC, CC-RIX, and DO across a range of QTL effect sizes and polygenic backgrounds for samples of 174 and 500 mice. The utility of replicate animals in the CC and CC-RIX for mapping QTLs rapidly decreased as traits became more polygenic. Only large sample populations of 500 DO mice were well-powered to detect smaller effect loci (7.5-10%) for highly complex traits (80% polygenic background). All results were generated with our R package musppr, which we developed to simulate data from these MPPs and evaluate genetic analyses from user-provided genotypes.

## Introduction

Multiparental populations (MPPs) are a powerful class of experimental cross for genetic studies of complex traits (de Koning and McIntyre 2017). Multiple isogenic founder strains were intercrossed to eventually produce offspring that possess recombinant genomes that encompass greater genetic diversity than traditional experimental crosses of two strains. MPPs have been developed across a wide range of model organisms, representing both animal models, such as heterogeneous stocks of mice and rats (Woods and Mott 2017), flies (Long et al. 2014), round worm (Noble et al. 2017), and yeast (Cubillos et al. 2013) as well as plants, including *Arabidopsis* (Kover et al. 2009), maize (Dell’Acqua et al. 2015), and rice (Bandillo et al. 2013). Here we focus on three related MPPs of the house mouse, *Mus musculus*: the Collaborative Cross (CC) panel of inbred strains (Collaborative Cross Consortium 2012; Srivastava et al. 2017), their F1 intercrosses (CC-RIX) (Threadgill et al. 2011; Schoenrock et al. 2018; Sun et al. 2021), and the Diversity Outbred (DO) population (Churchill et al. 2012).

The CC, CC-RIX, and DO share the same eight inbred founder strains (short names in parantheses): A/J (AJ), C57BL/6J (B6), 129S1/SvImJ (129), NOD/ShiLtJ (NOD), NZO/HlLtJ (NZO), CAST/EiJ (CAST), PWK/PhJ (PWK), and WSB/EiJ (WSB), which include both traditional laboratory and wild-derived strains and represent three subspecies of *Mus musculus* (Yang et al. 2007, 2011). As recombinant populations, the CC and DO can be used to map quantitative trait loci (QTLs) and have thus been used to genetically dissect a wide range of phenotypes. Traits that have been studied in the CC include behavioral traits (Philip *et al*. 2011), hematology traits (Kelada et al. 2012), airway damage (Tovar *et al*. 2022) and allergies (Kelada et al. 2014), susceptibility to influenza (Ferris et al. 2013) and SARS-CoV (Gralinski et al. 2015; Schäfer et al. 2022), homeostatic immune regulation (Hampton et al. 2021), and drug response (Mosedale et al. 2017, 2019, 2021); in the DO, serum cholesterol (Svenson et al. 2012), insulin secretion (Keller et al. 2019), glutathione metabolism (Gould et al. 2021), response to benzene exposure (French et al. 2015), bone and skeletal traits (Al-Barghouthi et al. 2021), and working (Ouellette et al. 2020) and short-term (Hsiao et al. 2020) memory.

The number of realized fully-inbred CC strains was lower than planned (The Complex Trait Consortium 2004), with approximately 70 strains yielded rather than 1,000 due to extinctions caused by allelic incompatibilities (Shorter et al. 2017). This reduction in the number of available strains, *i*.*e*., the number of unique genomes, reduced the potential power of genetic mapping studies in the CC; however, the inbred nature of the CC enables the use of replicates within and across experiments. Strain replicates can improve mapping power by reducing variation due to noise, and capture and identify strain-specific genetic effects and phenotypes, which can be caused by strain-specific variants and/or unique combinations of alleles across multiple loci. CC strains with unique phenotypes have been identified for a number of traits and diseases, including spontaneous colitis (Rogala et al. 2014), peanut allergy (Orgel et al. 2019), immune cell diversity (Dupont *et al*. 2021), and susceptibility to virally-induced neurological phenotypes (Eldridge et al. 2020), tuberculosis, (Smith et al. 2019) and *Salmonella* (Zhang et al. 2019; Scoggin et al. 2022).

Recently genomic studies have been a growing area of research for both the CC and DO, leveraging their genetic diversity in the presence of large genetic effects on molecular, *i*.*e*., omic traits, such as gene expression (Aylor et al. 2011; Keller et al. 2018) and chromatin accessibility (Keele et al. 2020), proteins (Chick et al. 2016) and their phosphorylation sites (Zhang et al. 2022), and lipids (Linke et al. 2020), across tissues and organs. The effects of biologically-related factors other than genetic variation, such as age, on gene expression and protein abundance have been studied in the DO mice (Takemon et al. 2021; Gerdes Gyuricza et al. 2022). Omic studies have also been performed in embryonic stem cells derived from DO mice (Skelly et al. 2020; Aydin et al. 2022). Genetic effects are strongly consistent between the CC and DO populations (Keele et al. 2021), supporting the use of both for joint analysis and validation of findings between populations.

QTL mapping power in the CC was initially evaluated prior to the development of the final panel of fully inbred strains, using far greater numbers of simulated genomes than were actually realized (Valdar et al. 2006). We updated mapping power estimates by simulating from readily available Collaborative Cross strains (Keele et al. 2019). Mapping power has also been assessed in the DO population (Gatti et al. 2014). Here we extend our previous approach of simulating from observed genomes, now across the CC, CC-RIX, and DO, to evaluate and compare the performance of commonly used genetic analyses: mapping QTLs and quantifying genetic architecture (in the form of estimating heritability). This work will aid researchers in designing and tailoring their experiments to maximize their efficiency for various goals (*e*.*g*., characterizing genetic architecture or mapping causal genetic variation), as well as provide recommendations for best practices in performing the quantitative and statistical analyses.

## Methods

### Sample populations

Generating realistic genetic data for MPPs from scratch poses multiple challenges. An ideal approach would involve full simulation of the various breeding designs from initial founder strains all the way to the reconstruction of founder haplotypes from marker genotypes in the final recombinant offspring. To avoid this complex and computationally intensive approach, we simulated data for CC, CC-RIX, and DO (Figure 1) from observed populations of CC and DO. This approach has the additional advantage of capturing the effects of genetic drift or any deviations from the breeding designs that occurred, better approximating real animals available to researchers.

**Figure 1.**
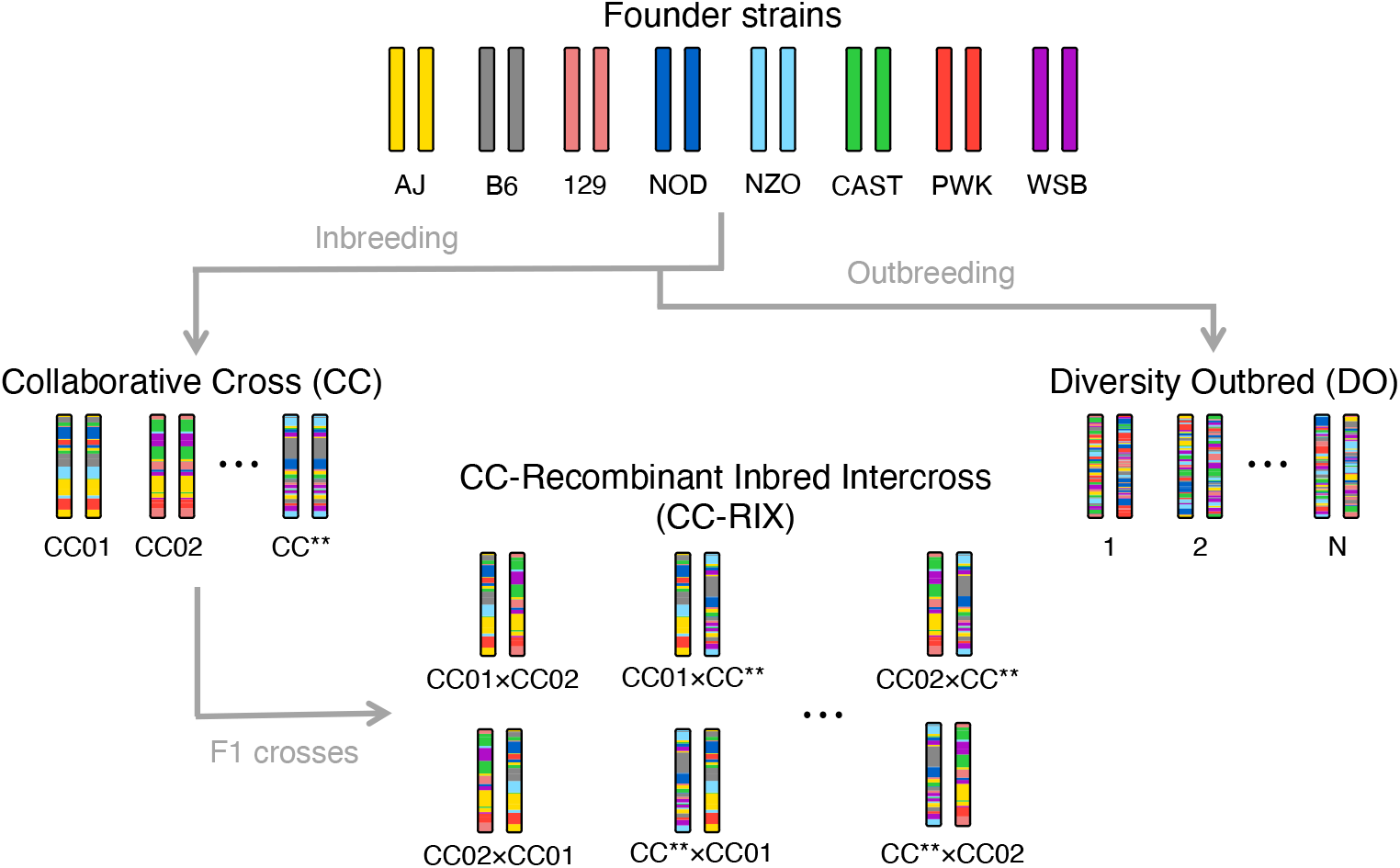
Cartoon diagram of the CC, CC-RIX, and DO populations. All the populations are recombinant and descended from the same eight inbred founder strains. Each genetically unique individual or strain is depicted as a single pair of chromosomes. For the CC-RIX, reciprocal F1s are depicted, *e*.*g*., CC01 × CC02 versus CC02 × CC01, indicating that the dam/sire identity of the CC parental strains are flipped. CC** represents the final CC strain. N is the total number of DO mice.

#### Genotype data

The CC sample population consisted of 116 mice (Keele et al. 2021) (female/male pairs from 58 strains) that were genotyped on an 11,000 marker array (MiniMUGA) (Sigmon *et al*. 2020). We used two DO sample populations, including 192 mice (Chick et al. 2016) genotyped on a 57,000 marker array (MegaMUGA) (Morgan and Welsh 2015) and 500 mice (Keller et al. 2018) genotyped on a 143,000 marker array (GigaMUGA). Founder haplotypes were probabilistically inferred using a hidden Markov model (HMM) implemented in the qtl2 R package (Broman *et al*. 2019). Genetic mapping in experimental crosses is commonly performed in terms of founder haplotypes rather than specific genetic variants, with uncertainty accounted for using a mixture model (Lander and Botstein 1989) (*i*.*e*., interval or linkage mapping) or regression-based approximations (Haley and Knott 1992; Martínez and Curnow 1992). Because the CC were genotyped on a sparser array than the DO, their founder haplotype reconstruction possesses greater uncertainty. To make these sample populations as comparable as possible in terms of quantified founder haplotype uncertainty and its effect on heritability estimation and QTL mapping, we imputed founder haplotype probabilities at the same 64,000 loci (*i*.*e*., pseudomarkers) spanning the genome and then imputed the founder haplotype pair (*i*.*e*., diplotype) based on greatest probability. We note that the genetic data and corresponding results reported here are idealized to some extent, and real data will be subject to genotype uncertainty. We derived the founder diplotypes for the 1,653 CC-RIX F1s, representing all possible pairings between 58 CC strains (ignoring parent-of-origin features, *e*.*g*., sex chromosomes and mitochondria). See Appendix A for greater detail on how diplotypes were processed for all sample populations.

#### CC-RIX F1 selection

For the CC-RIX, given that it is unlikely researchers would collect data for a full set of all possible F1s (1,653 for 58 CC parental strains), we evaluated three classes of simulated populations. These three types of CC-RIX populations are not meant to represent all possible approaches to selecting F1s and designing a CC-RIX experiment, though they do possess distinct features described below.

For the first class of CC-RIX population, F1s were selected such that each CC strain is a parent for two F1s, as is possible with a rotational breeding scheme (*e*.*g*., CC001 × CC058, CC001 × CC002, CC002 × CC003, …, CC057 × CC058). We refer to these populations as “balanced” because all CC strains are observed equally as parental strains. Each F1 was simulated with multiple replicates. Balanced samples of CC-RIX F1s possess the same overall allele frequencies as a corresponding sample of the parental CC strains, allowing us to decouple the effects of allele frequency from how the CC-RIX population structure and overall heterozygosity affects heritability estimation and mapping power.

We also evaluated simulated populations that were composed of randomly sampled CC-RIX F1s, which we refer to as “unbalanced”. This results in unequal representation of parental CC strains by their offspring F1s in the resulting CC-RIX sample population. We evaluated unbalanced CC-RIX F1 sets in CC-RIX-only populations and in combination with the parental CC strains. Examples of each class of CC-RIX sample populations are shown in Figure S1.

### Heritability

Heritability, the proportion of variation in a population explained by genetic relationship, is commonly used to assess evidence that a phenotype is genetically controlled. It is important to note that heritability is specific to a population, and thus cannot be extrapolated across populations. Furthermore, it can be influenced by other sources of variation in the data, such as the measurement error of the phenotype, which will likely reduce heritability estimates.

Depending on the sample population, heritability can be estimated using different approaches and even decomposed into multiple components, such as the proportion of variation explained by all genetic effects, *i*.*e*., broad-sense heritability (commonly symbolized as *H*^2^) and the proportion of variation explained by additive genetic effects, *i*.*e*., narrow-sense heritability (*h*^2^) (Lynch and Walsh 1998). In this work, we use the *h*^2^ notation more generally and will include subscripts to specify components of heritability in certain contexts. For studies of inbred strains with replicates, as is possible with the CC, an intraclass correlation can be used (as in Yam *et al*. 2021), though it is less appropriate in the outbred CC-RIX and not possible in the DO.

#### Heritability model

Linear mixed effects models (LMMs) offer an appealing general approach that can be adjusted for each of the CC, CC-RIX, and DO. We estimate heritability using the following general LMM:

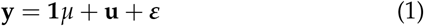

where 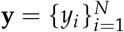 is the phenotype vector for a sample population of *N* mice, *µ* is the intercept, and **u** and ***ε*** are *N*-length random vectors. **u** can be referred to as the polygenic effect, here representing structured error (*i*.*e*., population structure) and is modeled as 𝒩 (**0, K***τ*^2^) where **K** is an *N× N* (often additive) genetic relationship matrix (*i*.*e*., kinship matrix) and *τ*^2^ is the corresponding variance component. ***ε*** is unstructured error, distributed according to 𝒩 (**0, I***σ*^2^) where **I** is the *N × N* identity matrix and *σ*^2^ is its variance component. Heritability is then calculated as the ratio of variation due to genetics to total variation:

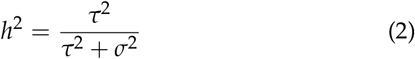

To obtain unbiased estimates of heritability, the variance component parameters are estimated through optimizing the restricted maximum likelihood (REML) (Patterson and Thompson 1971). The heritability LMM can be fit with various software packages; here we compare results from the qtl2 R package (Broman et al. 2019), our miQTL R package (available at https://github.com/gkeele/miqtl), and the sommer R package (Covarrubias-Pazaran 2016).

For sample populations that include strain or F1 replicates, such as in the CC and CC-RIX, the kinship matrix can be characterized as **K** = **ZK**_*M*_ **Z**^T^ where **Z** is the *N× M* strain/F1 identity matrix that maps the *N* individuals to *M* strains/F1s (*M* < *N*) and **K**_*M*_ is the *M ×M* kinship matrix encoding the overall genetic relationship between the *M* strains/F1s.

#### Expanded heritability model with non-additive component for replicates

If replicates are included in the sample population, Eq 1 can be expanded to include two components of heritability:

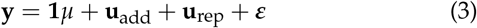

where **u**_add_ is equivalent to **u** from Eq 1 in the presence of replicates 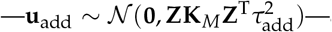and **u**_rep_ is a random effect specific to strain/ 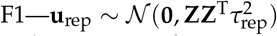 The two components of heritability are then estimated as

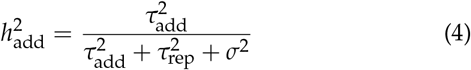

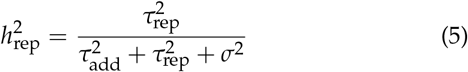

where 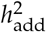 is the proportion of variation explained by additive genetic effects, *i*.*e*., narrow-sense heritability, and 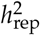 is the proportion of variation explained by strain/F1 identity, such as due to epistasis between loci with combinations of alleles unique to a given strain/F1. From the three software packages we used, only sommer allows for flexible specification of multiple random effects, each with their own covariance matrix.

#### Kinship matrix for heritability

The true kinship matrix **K** is unknown and must be estimated from genotypes (haplotypes or SNPs), pedigree information, or both (Cheng et al. 2013). Here our focus is not a thorough analysis of how to best estimate the kinship matrix, though we do evaluate a number of options. For a SNP-based kinship matrix, we use the form described in Endelman and Jannink (2012) with selection parameter set to 0. See Appendix A for details on imputing SNPs from the founder haplotypes. For a haplotype-based kinship matrix, we used qtl2 (Broman et al. 2019), which calculates 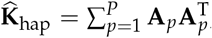 **A**_*p*_ is the scaled founder haplotype dosage matrix at locus *p*. Recently, Feldmann *et al*. (2022) proposed the average semivariance (ASV) transformation for kinship matrices, which resulted in improved heritability estimation. Here we also evaluated ASV forms of each kinship matrix as 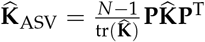, where 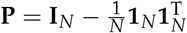 is the mean-centering matrix.

#### Heritability simulations

We adapted our approach used in Keele *et al*. (2019) to simulate phenotype data with specified heritability; see Appendix B for greater detail. When simulating a single additive component of heritability for a given population, we simulated 1,000 data sets for each specified level of heritability. Heritability was evaluated across a grid, ranging from 0 to 1 with increments of 0.05. Heritability was then estimated using one of qtl2, miQTL, or sommer, and summarized with the mean and the 95% estimate interval (defined by the 0.025 and 0.975 empirical quantiles). Within this simulation framework, we also compared using kinship matrices estimated from SNPs and founder haplotypes, as well as ASV forms of both.

We extended the previous simulation approach for heritability with two components by fixing the total heritability at a specified level (*e*.*g*., 90%) and varied the ratio 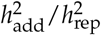 across a grid, ranging from 0 to 1 with increments of 0.1. Only 100 data sets were simulated per parameter setting because fitting the two component heritability model in sommer is more computationally intensive and thus slower to fit. Means and 95% estimate intervals were used as summaries as before, but now for each component of heritability separately as well as their sum.

We performed various comparisons of heritability estimation across the sample populations of CC, DO, and balanced and unbalanced CC-RIX by generating simulated populations derived from their genomes. Generally we down sampled mice to make the number of total mice consistent across populations for a given comparison. For example, we randomly sampled 174 DO mice from the sample population of 192 to compare to the 174 CC mice (three per 58 strains). When evaluating two components of heritability, we simulated larger populations of 522 mice given that each genome must be observed with multiple replicates to distinguish the strain- or F1-specific component from the additive one.

### QTL mapping

The chromosomes of individuals from MPPs are mosaics of the founder haplotypes formed through recombination events that occurred during outbreeding (Figure 1). This mixing process randomizes loci from each other, ignoring linkage disequilibrium (LD) and population structure, allowing genetic variation at loci to be causally associated with traits. We evaluate mapping power in the context of conventional single locus genetic mapping, though note that these approaches could be extended to multi-locus models and epistasis.

#### QTL model

The underlying model used during QTL mapping is similar to Eq 1 for heritability, however a locus effect is evaluated at loci spanning the genome (*i*.*e*., a genome scan):

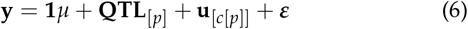

where **QTL**_[*p*]_ = **A**_*p*_ ***β***_QTL_ is the effect of locus *p*, **A**_*p*_ is the *N×* 8 scaled founder haplotype dosage matrix at locus *p* (assuming an additive model), ***β***_QTL_ is the 8-vector of founder haplotype effects, **u**_[*c*[*p*]]_ is the random polygenic effect vector with respect to chromosome *c* (on which locus *p* is located), and all other terms as previously defined. This represents a leave-one-chromosome-out (LOCO) approach (Yang et al. 2014; Gatti et al. 2014) in which **u**_[*c*[*p*]]_ ∼ 𝒩 (**0, K**_[*− c*[*p*]]_), meaning **K** is estimated from all markers excluding those on locus *p*’s chromosome, which avoids any of the effect of locus *p* from being absorbed into the random term and increases mapping power. We used qtl2 to perform all genome scans.

#### QTL significance thresholds

Instead of using permutations (Churchill and Doerge 1994) to estimate significance thresholds that control the genome-wide family-wise error rate (FWER), we used parametric bootstrap samples generated from the null model (Eq 6 excluding the QTL term). We note that the bootstrap samples are generated given the true value of heritability, which would normally need to be estimated from the data and thus subject to error. Our goal is to produce thresholds that can be used across simulated data for a given population with the same parameter settings. Thresholds were calculated as quantiles from extreme value distributions fit from the maximum LOD scores across the null bootstrap samples (Dudbridge and Koeleman 2004). For a QTL to be correctly detected, we required its peak marker to be located within 1.5 LOD support intervals (Dupuis and Siegmund 1999), which we discuss in great detail in the next section of the Methods.

In the context of an omic trait, such as a gene’s expression, there is strong biological support for genetic variants that are nearby the trait’s genomic positions (*i*.*e*., local) having strong genetic effects, such as *cis*-eQTLs. This prior evidence allows for more lenient thresholds to be used to more powerfully detect local QTLs (Keele et al. 2020). Here we use a simple approach to lenient thresholds by repeating the previous genome-wide procedure, but now reducing the multiple testing burden to the loci on the traits’s chromosome (*i*.*e*., local chromosome). This is analogous to only considering the chromosome on which a gene is encoded for *cis*-eQTLs.

#### Confidence interval for QTL location

The goal of QTL mapping is often to identify candidate genes or functional variants that underlie a QTL. But just as estimation of the effects of the QTL are subject to error, so is the estimation of QTL location. Furthermore, most often the causal genetic variant itself is not being tested as a QTL, but rather a locus that is in LD with it. This is particularly relevant in MPPs, where it is conventional to initially detect QTL using sparser scans based on founder haplotypes at intervals across the genome (Eq 6). A confidence or support interval for the location of QTL summarizes this uncertainty and can prioritize a genomic region for specific candidate genes or genetic variants.

Approximate likelihood-based support intervals are commonly used due to ease of computation, which were statistically characterized for F2 intercrosses and backcrosses (Dupuis and Siegmund 1999) but not for MPPs such as the CC, CC-RIX, or DO. Empirical sampling-based approaches have also been proposed, such as non-parametric bootstrapping (Visscher et al. 1996), though have been found to perform poorly with sparse markers (Manichaikul et al. 2006). Here we evaluate approaches for estimating QTL intervals in CC, CC-RIX, and DO populations, including likelihood-based—LOD support intervals and approximate Bayes credible intervals (Broman and Sen 2009)—and sampling-based intervals— parametric bootstrap, parametric permutation, and Bayesian bootstrap (Rubin 1981). See Appendix C for more details on each method.

#### QTL simulations

We extended our approach to simulating mapping data in the CC (Keele *et al*. 2019) to the CC-RIX and DO populations. We evaluated QTL mapping performance across the populations, comparing power, mapping resolution, and interval summaries for QTL location, while varying the number of mice and the proportion of phenotypic variation due to a QTL and the cumulative effect of background genetic loci (approximated through a polygenic effect). All combinations of QTL effect size ranging from 10% to 40% in increments of 5% and polygenic background of 5% (essentially a Mendelian trait), 30%, and 55% were performed for CC, CC-RIX, and DO populations of 174 and 500 mice. We also considered the context of small QTLs (<10%) in highly complex traits (polygenic background of 80%). For this case, we simulated QTL effect size of 1%, 2.5%, 5%, 7.5%, and 10% for sample populations of 174 and 500 mice.

For CC and balanced CC-RIX samples of 174, three replicates per 58 strains/F1s were simulated. We used 174 because we had genetic data for 58 CC strains and a smaller DO sample population of 192 mice. Using three replicates per CC strain resulted in 174 mice, whereas four replicates (232 mice) would have exceeded the smaller DO sample. For samples of 500, ten replicates per 50 strains/F1s were simulated for CC and balanced CC-RIX. Unbalanced CC-RIX populations did not include any replicates.

Across all simulation scenarios (QTL effect size, polygenic background, population, sample size), we randomly selected 1,000 loci as QTLs from which to simulate data. The variation explained by the QTL and polygenic background were strictly controlled with respect to the sample population. Each QTL was simulated as having a bi-allelic effect evenly split across the founder strains. We discuss the implications of this further in the Results and Discussion.

We also performed reduced sets of simulations to compare mapping power based on CC strain means to individual-level data as well evaluate how differences in allele frequency and heterozygosity across the CC, CC-RIX, and DO populations may affect mapping results. For the comparison of CC strain means to individual-level data, 1,000 QTLs were simulated in CC populations of 116 mice (2 replicates per 58 strains) across the previously used low-to-moderate polygenic backgrounds (5%, 30%, and 55%). To evaluate the effect of allele frequency and heterozygosity, we scaled the QTL effect size with respect to a reference population, one that is fully inbred with perfectly balanced allele frequencies. We simulated 1,000 QTLs in populations of 174 mice with a polygenic background of 30%. For both reduced analyses, we varied QTL effect size from 10% to 40% in increments of 5%. See Appendix D for further explanation of our simulation approach for QTL data.

## Results and Discussion

We simulated data using previously observed genomes of CC, CC-RIX, and DO mice to evaluate heritability estimation and QTL mapping; see Methods for more details on the populations, the simulations, and statistical methods used.

### Heritability estimation performance

#### ASV transformation improves accuracy and precision

We first compared heritability estimation across kinship matrices (haplotype-based and SNP-based, both non-ASV and ASV forms) and statistical software (qtl2, miQTL, and sommer) that can fit the same LMM (Figure S2). Haplotype-based and SNP-based kinship matrices with matching ASV status performed similarly, which is not surprising given that the SNPs were imputed from the haplotypes. This does confirm that these estimators produce similar genetic relationship matrices. In terms of software packages, miQTL and sommer produce essentially identical estimates of heritability, performing best with ASV kinship matrices. In contrast, qtl2 performs best in the DO with non-ASV haplotype-based matrices and is negatively biased when estimating heritability in the CC. For these reasons, we present the remaining heritability results using miQTL, which is computationally faster than sommer, and using ASV haplotype-based kinship matrices.

#### Replicates improve precision

We first evaluated heritability estimation in small samples of 50 and 100 mice in the CC and DO (Figure 2). For a sample of 50 mice, the mean heritability across the simulations was more biased at the tails of the distribution. More importantly, the 95% estimate intervals cover the full support of heritability; in other words, regardless of the true value of heritability, the interval ranges from 0% to 100% (Figure 2A-B). Doubling the number of animals to 100 increases the precision of heritability estimation (Figure 2C-D), most notably in the CC population with two replicates per strain. For a 50% heritable trait in 100 DO mice, the 95% estimate interval still ranges from 0% to 100%, whereas in the CC, it ranges from 30% to 65%. In the CC, there was also greater variability in heritability estimates when the true heritability was lower, though the widest intervals are still notably narrower than in the DO sample population of the same number of mice.

**Figure 2.**
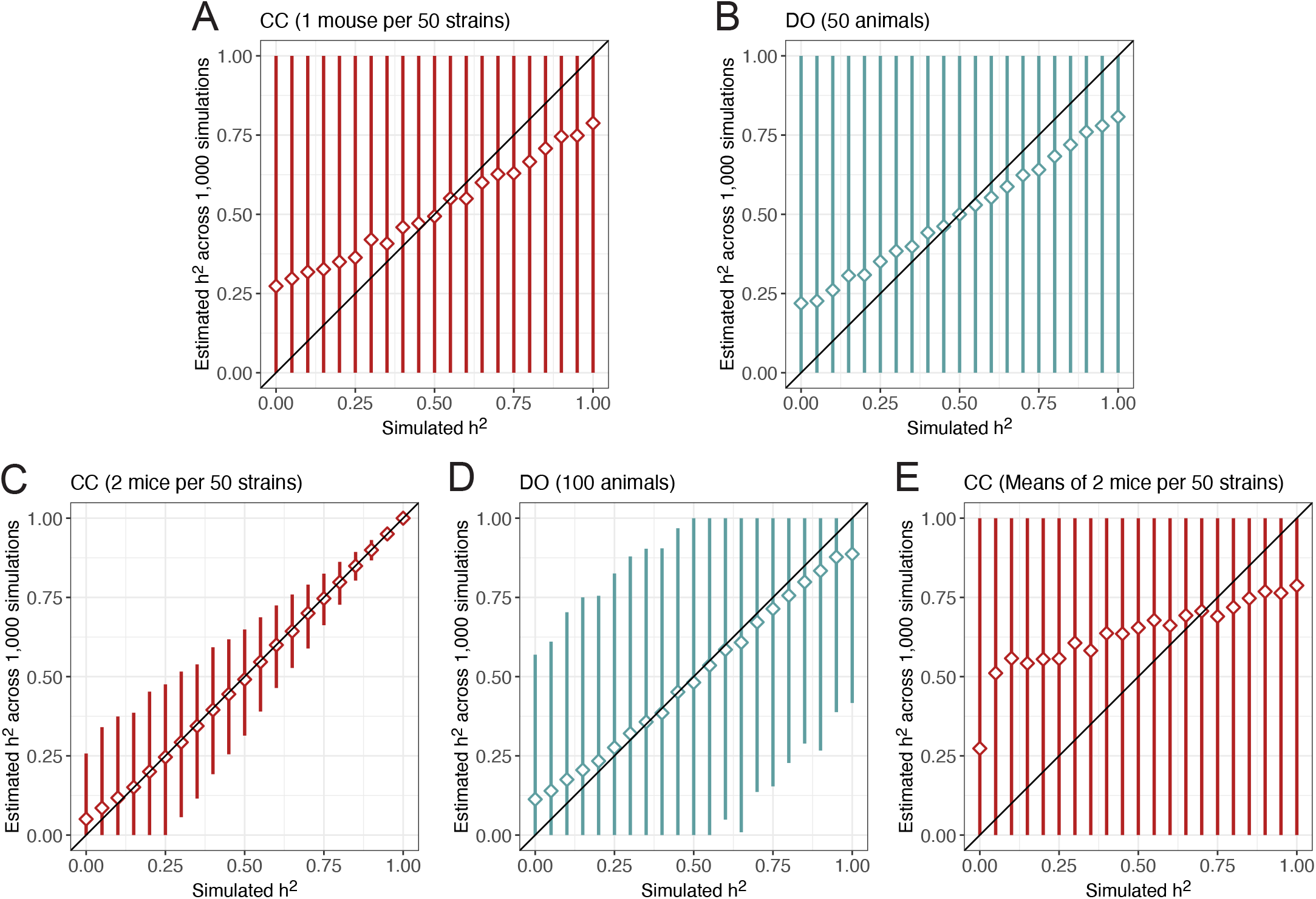
Performance of heritability estimation in data simulated for (A) one mouse per 50 CC strains, (B) 50 DO mice, (C) two mice per 50 CC strains, (D) 100 DO mice, and (E) strain means from two mice per 50 CC strains. Diamonds represent the mean estimated heritability across 1,000 simulations from the true heritability. Vertical line segments represent middle 95% intervals across the 1,000 simulations. Black diagonal lines indicating the heritability estimate is equal to the true value (y = x) included for reference.

We also evaluated heritability estimation using CC strain means (Figure 2E), which are commonly used for QTL mapping, and observed upward bias and intervals that cover the full support of heritability. The upward bias is consistent with the expected reduction in noise on a mean, and the wide intervals are due to the loss of the within-strain correlation information. The underlying heritability model could be adjusted for strain means, incorporating the number of replicates per strain and standard errors on the strain means, which would likely correct bias and increase precision. However, this would require statistical models tailored for the data with replicates. If this is not the case, this finding suggests that replicates should not be reduced to strain-level summaries for the purpose of estimating heritability.

We expanded our evaluation of heritability estimation to include three classes of CC-RIX populations while fixing the number of total mice at 174 (three replicates per 58 strains/F1s for CC and Balanced CC-RIX; Figure 3) and 500 (ten replicates per 50 strains/F1s for CC and Balanced CC-RIX; Figure S3). Compared to populations of 50 or 100 mice, larger populations of 174 or 500 are able to estimate heritability more accurately and precisely. To more directly compare how the number of replicates versus the total number of mice influence the accuracy and precision of heritability estimation across the populations, we performed 1,000 simulations for varying numbers of replicates and total mice at two levels of heritability (25% and 75%) (Figure 4). Overall, bias was not an issue for any of the populations except for small samples of CC (50 mice with no replicates) and DO (50-100 mice) (Figure 4A). Precision of heritability, however, was notably improved through replicates rather than total number of mice—a sample of five replicates per 10 CC strains (50 mice total) has similar or better precision than a DO population of 400-500 mice (Figure 4B). For example, the 95% estimate interval width in a sample of five mice from 10 CC strains is 15.1% compared to 100.0% in 100 DO mice, 43.3% in 400 DO mice, and 32.7% in 500 DO mice when the true heritability is 75%.

**Figure 3.**
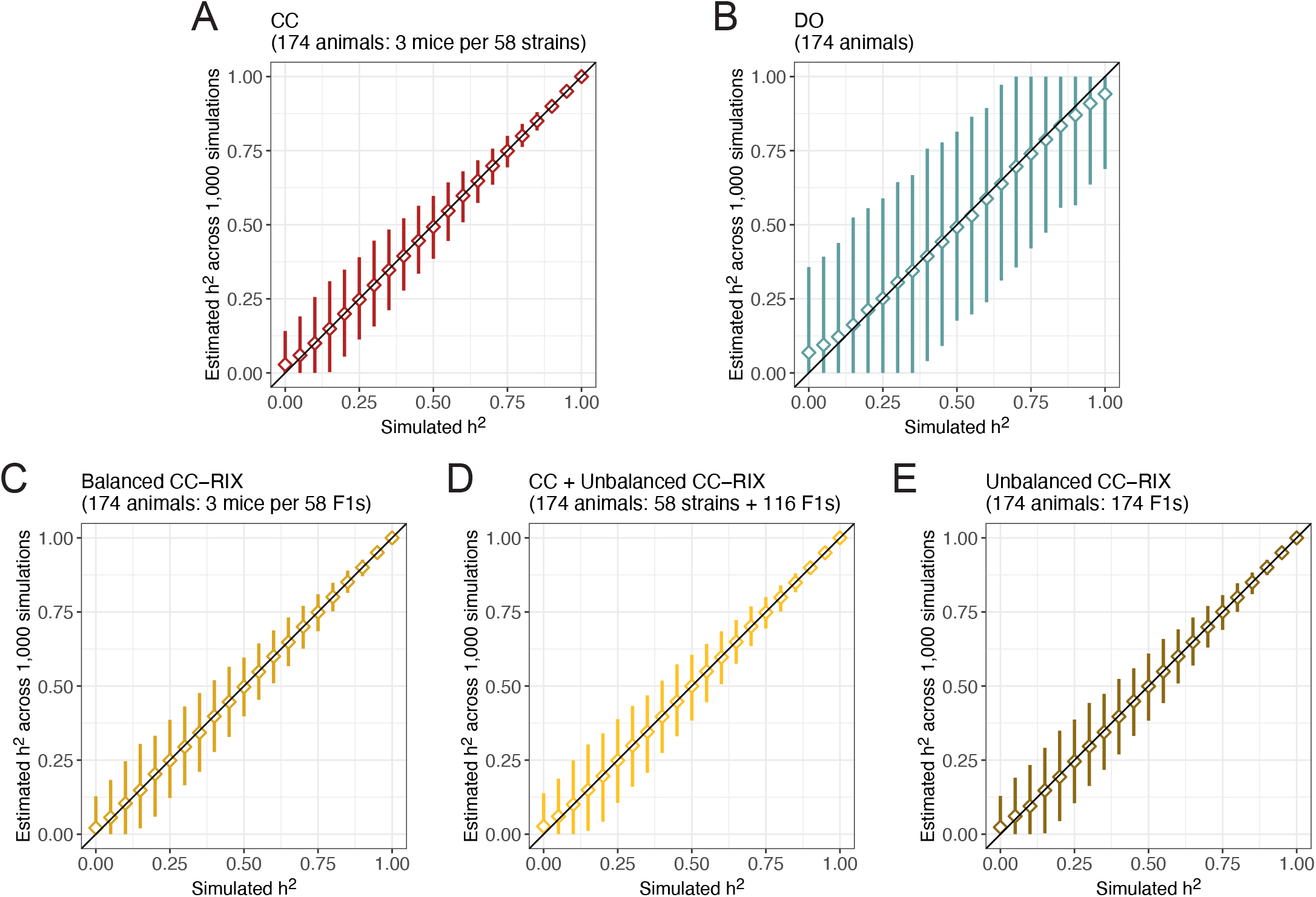
Performance of heritability estimation in data simulated for 174 mice from (A) CC, (B) DO, and (C-E) CC-RIX populations. Diamonds represent the mean estimated heritability across 1,000 simulations from a given true heritability. Vertical line segments represent middle 95% intervals across the 1,000 simulations. Black diagonal lines indicating the heritability estimate is equal to the true value (y = x) included for reference. See Figure S3 for results from simulations of 500 mice.

**Figure 4.**
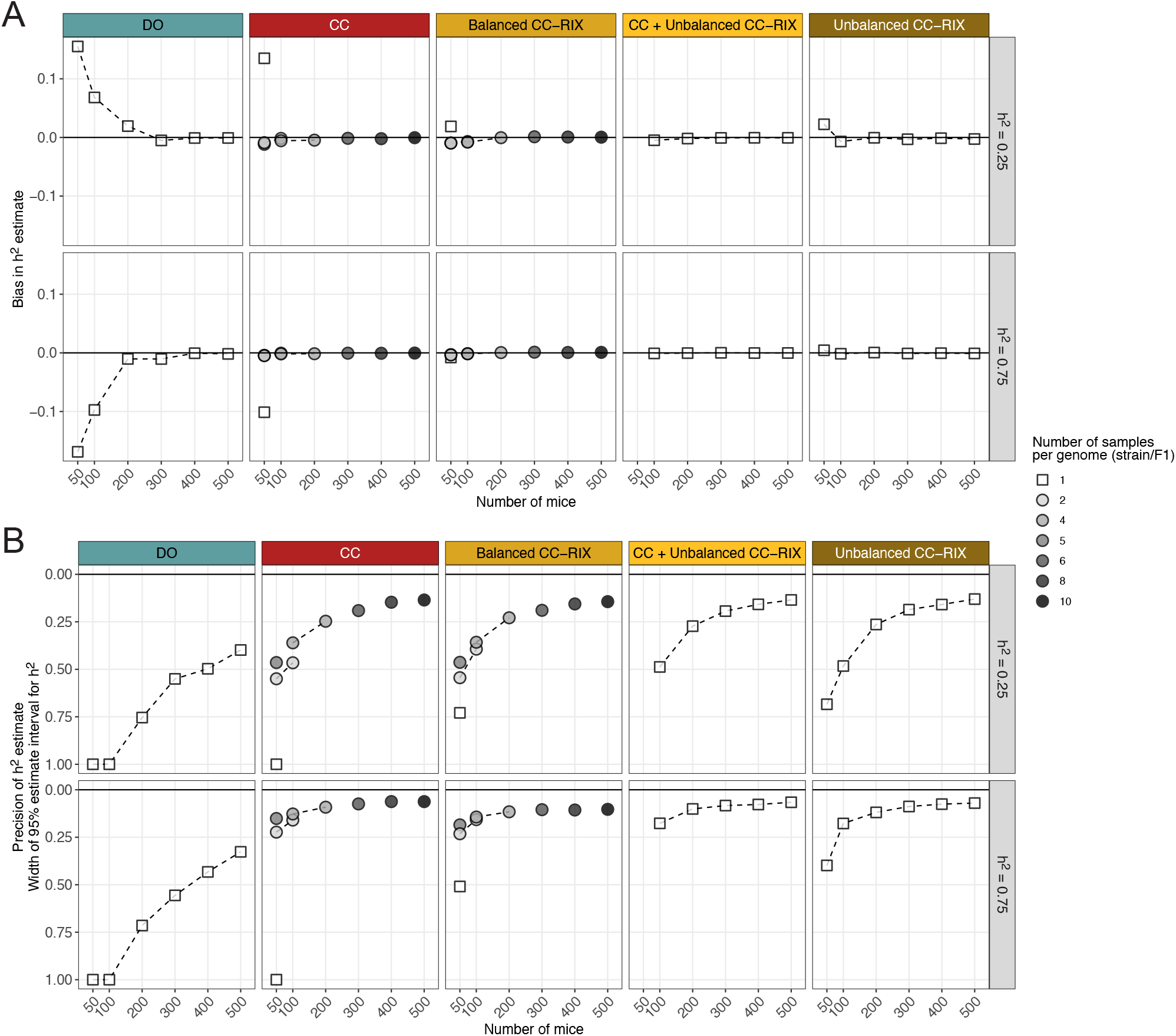
Comparison of (A) bias and (B) precision in heritability estimation across CC, CC-RIX, and DO populations, number of replicates, and total sample size. Points represent summaries over 1,000 simulations for a given population type (columns) and true heritability (rows). The number of replicates per genome is indicated by point shape and grayscale. Bias and precision are reported as the mean and 95% estimate interval width across the 1,000 simulations, respectively. Black horizontal lines indicating no bias and perfect precision (y = 0) are included for reference. Dashed lines connect summaries with the same number of samples per genome.

The CC and CC-RIX populations provide more precise heritability estimation than the DO because they possess greater variation in inter-relatedness across their sample populations (essentially, population structure) (Figure S4) in the form of genetic replicates and/or shared parental CC strains (for the CC-RIX). In data simulated for 174 mice and heritability set to 80%, all populations produce low bias, but the 95% estimate interval width for the DO is greater than 45% and around 10% in the CC and CC-RIX (Figure S4B). Notably, population structure also has implications for QTL mapping power. For example, not accounting for population structure in QTL mapping, produces many false positives, including many with strong associations (LOD score > 10), in the unbalanced CC-RIX populations (Figure S4C). This emphasizes the importance of accounting for population structure though the LMM in QTL mapping. We will discuss QTL mapping power in greater detail further on.

#### Replicates of CC-RIX distinguish additive and non-additive components of heritability

We next performed simulations with two components of heritability: an additive component as before that reflects the additive kinship matrix, as well as a strain- or F1-specific component. Because replicate DO samples are not observable, we only evaluated CC and CC-RIX populations (Figure 5). All simulations had a cumulative heritability of 90% in populations of 522 mice. Across 100 simulations in the CC, the ratio of estimated components are generally unbiased but with 95% intervals ranging from 0% to 90% whereas the cumulative heritability was accurately and precisely estimated at 90% (Figure 5B). In the CC, it is unsurprising that cumulative heritability can be accurately estimated whereas the individuals components are indistinguishable given that the strains of the CC are largely equally related and thus the majority of population structure in a sample population comes from replicates, confounding the additive and strain-specific effects. This is further confirmed by how fitting an additive-only heritability model to data simulated with both additive and strain components generally returns the cumulative heritability total rather than the additive component (Figure S5A).

**Figure 5.**
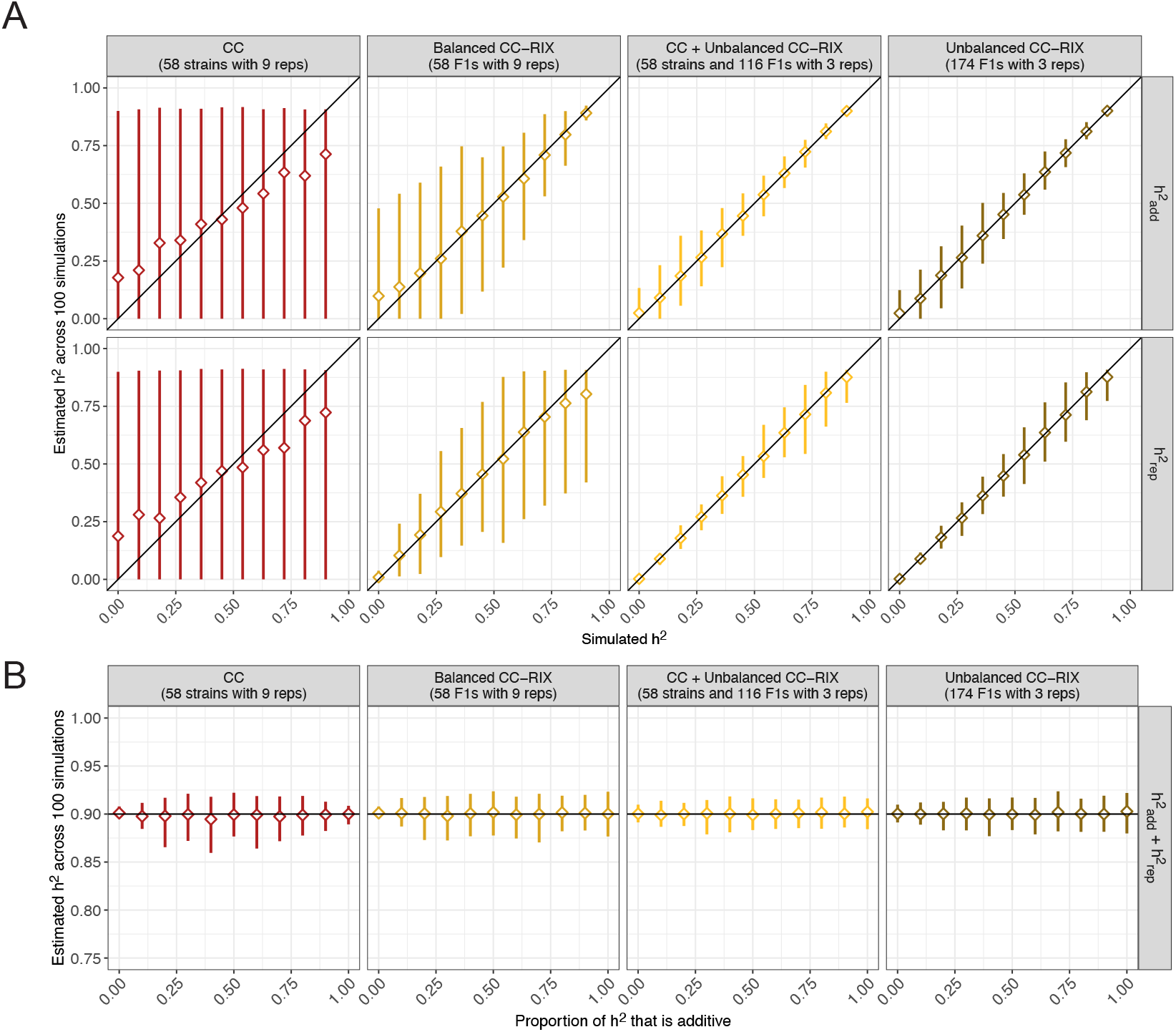
Performance of two component heritability estimation in data simulated for 522 mice from CC and CC-RIX populations. The sum of both components was fixed at 90%. Heritability was estimated either in terms of (A) each component, with the additive component 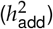 shown in the top row and the strain- or F1-specific component 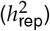 shown in the bottom row, or as (B) the sum total. Diamonds represent the mean estimated heritability across 100 simulations from the true heritability. Vertical line segments represent middle 95% intervals across the 100 simulations. Black diagonal lines indicating the heritability component estimate is equal to the true value (y = x) included for reference. Black horizontal lines indicating the true value of the sum of heritability components (y = 0.9) included for reference.

All CC-RIX populations better distinguish the additive and non-additive components of heritability than the CC, particularly the unbalanced CC-RIX populations (Figure S1). Fitting an additive-only heritability model in simulated unbalanced CC-RIX data with two components is also better able to capture only the additive component, though some of the non-additive variation does appear to upwardly bias the estimate (Figure S5B). These findings demonstrate the utility of the CC-RIX and their unique half-sibling F1s for distinguishing additive and non-additive genetic effects.

### QTL mapping power curves for traits with low-to-moderate polygenic backgrounds

We evaluated the power to detect QTLs with effect sizes ranging from 10% to 40% by 5% increments in traits with low-to-moderate polygenic backgrounds (5%, 30%, and 55%) (Figure 6A). This range of genetic effect parameters covers a phenotype spectrum ranging from simple Mendelian traits to more genetically complicated ones. Sample populations of 174 (three replicates per 58 strains/F1s in CC and balanced CC-RIX) and 500 (10 replicates per 50 strains/F1s) were simulated and evaluated.

**Figure 6.**
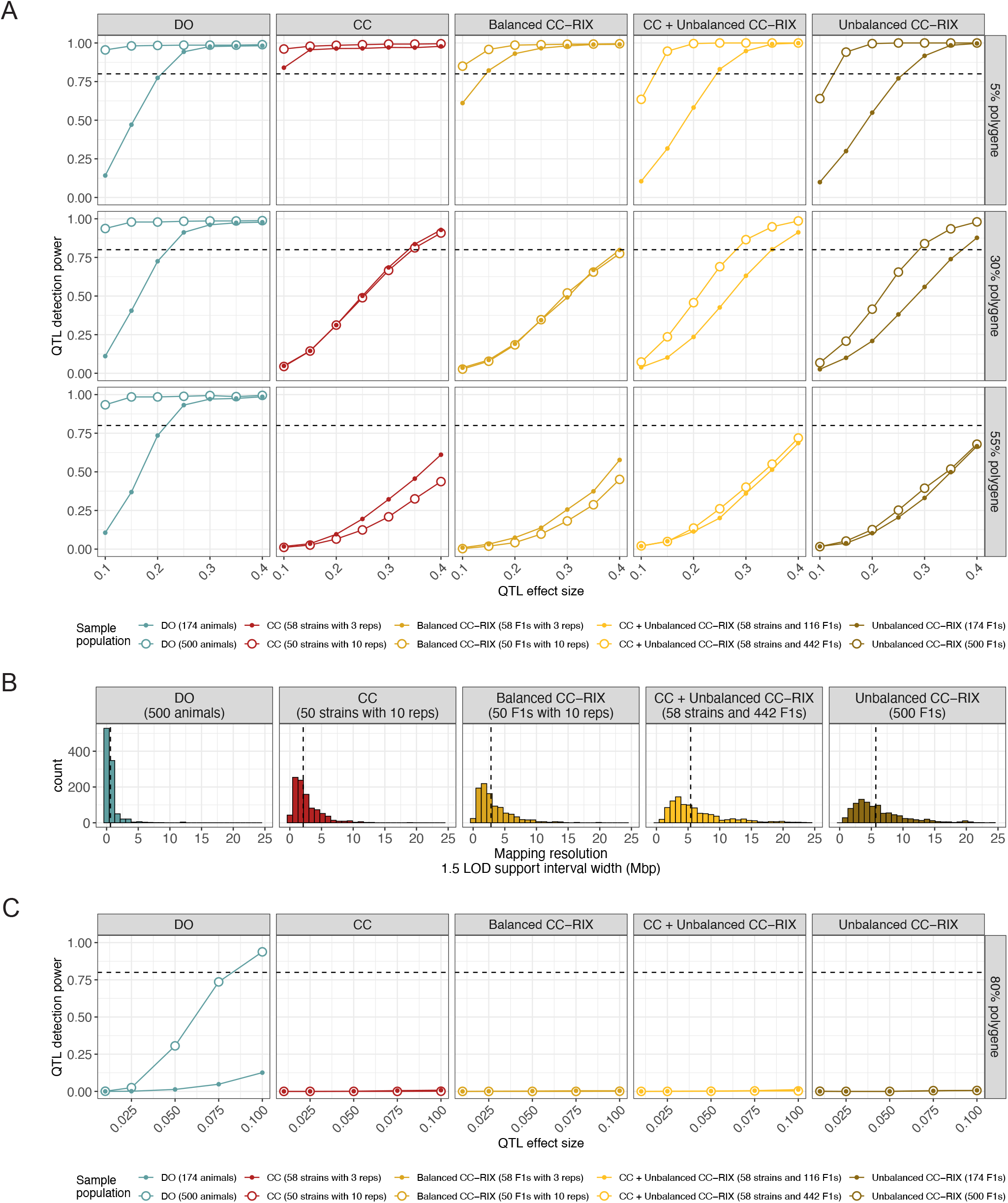
Performance of QTL mapping in data simulated for CC, CC-RIX, and DO populations based on genome-wide significance. (A) QTL power curves across populations (columns) and low-to-moderate polygenic backgrounds (rows). Sample populations were composed of 174 (solid points) and 500 mice (open points). Horizontal dashed lines at 80% power included for reference. (B) Histograms of the 1.5 LOD support interval width for simulations of a 40% QTL with 5% polygenic background across the populations (columns). Vertical dashed lines represent the median for each population. (C) QTL power curves across populations (columns) in a highly polygenic background (80%). Sample populations were composed of 174 (solid points) and 500 mice (open points). Horizontal dashed lines at 80% power included for reference. For power estimates based on lenient local chromosome-wide FWER control, see Figure 7.

#### Population structure reduces QTL mapping power

For nearly monogenic traits (5% polygenic background), all populations were well-powered to detect QTL with effect sizes within the range of 10% to 40% (Figure 6A [top row]). The value of replicates for QTL mapping is maximized for traits with low polygenic effect, with the CC and balanced CC-RIX mostly out-performing even the DO. As the genetic background of the trait became more polygenic, populations with high levels of structure (*i*.*e*., unequal relatedness across individuals) due to replicates or CC-RIX half-sibling F1s performed more poorly. With a 30% polygenic effect, populations of 174 with replicates (CC and balanced CC-RIX) were essentially equally powered as populations of 500 (Figure 6A [middle row]). This becomes more pronounced with 55% polygenic traits; populations of 174 with replicates were better powered than corresponding populations of 500 because they included more distinct genomes (58 CC strains compared to 50) (Figure 6A [bottom row]). The unbalanced CC-RIX populations of 500 are only slightly better powered than the corresponding populations of 174. In contrast to the CC and CC-RIX populations, mapping power in the DO was highly consistent across background polygenicity.

#### DO provide narrower QTL intervals than CC or CC-RIX

Mapping in the DO produced narrower QTL intervals, as measured by LOD support intervals, followed by CC and balanced CC-RIX, and finally the unbalanced CC-RIX populations (Figure 6B). The QTL interval width is expected to be inversely related to the number of distinct recombination events that are observed in a mapping population. These results are consistent with this expectation, given that the DO possess far more recombination events that occurred during additional outbreeding generations over the CC and CC-RIX, as well as possessing more genetically unique individuals than populations with replicates. The unbalanced CC-RIX populations potentially include the fewest recombination events because not all the available 58 CC strains are necessarily selected as parental strains, resulting in wider QTL intervals. We also note that the DO continue to be intercrossed to produce new generation, with each subsequent generation accruing more recombinations and thus finer mapping resolution in principle. LOD support intervals were used as a summary of mapping resolution due to ease of computation. Further on we more deeply evaluate the statistical performance of QTL interval estimates.

#### Individual-level data improved QTL mapping power over strain/F1 means

In the CC, it has been common practice to perform QTL mapping based on strain-level summaries (*e*.*g*., means) rather than individual mouse-level data. Often specific animals are not even genotyped, and instead resource genotypes summarized from multiple ancestors are used (available at http://csbio.unc.edu/CCstatus/index.py?run=FounderProbs). Strain summaries also have the added benefit of reducing the data and the resulting computational burden. We sought to evaluate how this strain-level approach compared to use of individual-level data in terms of QTL mapping power. Based on simulations of 1,000 QTLs in 116 mice (two replicates per strain), we observed reduced mapping power with CC strain means compared to individual-level data. For a 40% QTL in a 30% polygenic background, we observed 88% power in CC strain means compared to 98% in CC individuals (Figure S6). The disparity in power varied with QTL effect size and polygenic background, but individual-level data universally performed better. We primarily report results using strain means (and F1 means for balanced CC-RIX) given this approach has been commonly used, but these findings suggest that the use of individual-level data results in appreciable gains in mapping power.

### QTL mapping power curves for traits with a highly polygenic background

We next evaluated power to detect QTLs with small effect sizes (1% to 10%) in traits that are highly polygenic (80%) (Figure 6C). These simulations are meant to approximate highly heritable traits that are controlled by many genetic loci with small individual effects, such as height in humans (Vinkhuyzen et al. 2013), which has heritability estimates around 80%. As before, sample sizes of 174 and 500 were evaluated.

#### Only large samples of DO are well-powered to detect small effect QTLs in a highly polygenic background

Only sample populations of 500 DO mice were potentially powered to detect QTLs in a highly polygenic background, whereas CC and CC-RIX populations had no power. Specifically, samples of 500 DO mice were well-powered (>80%) to detect QTLs that explain 7.5% to 10% of phenotypic variation. A key takeaway is that as the genetic background becomes more polygenic, the value of a genetic replicate in terms of mapping power decreases. Though a genetic replicate will still greatly increase the precision of heritability estimates, they contribute essentially no improvement to mapping power for traits with polygenic backgrounds of 30% or greater.

### QTL mapping power curves for omic traits using lenient local significance thresholds

Genetic variants that strongly affect an omic trait are often in close proximity to the trait’s genomic coordinates, such as a *cis*-eQTL that affects the availability of a gene’s promoter for the initiation of transcription. In this context, analyses focused on the local genomic region of a trait can improve power by reducing the multiple testing burden (Keele et al. 2020). We re-evaluated the power to detect QTLs in the prior scenarios, but now assuming all 1,000 simulated QTLs are local and now restricting our testing to the local chromosome of each trait (Figure 7). Detecting QTLs based on local significance improves power, most notably in the CC and CC-RIX for traits with low-to-moderate polygenic backgrounds, suggesting that they have more utility for mapping local QTLs for omic studies than traditional complex traits, in which they are often under-powered (Figure 6). In a highly polygenic background, the power gains are less noticeable and the CC and CC-RIX do not exceed 12% power for any QTL effect size.

**Figure 7.**
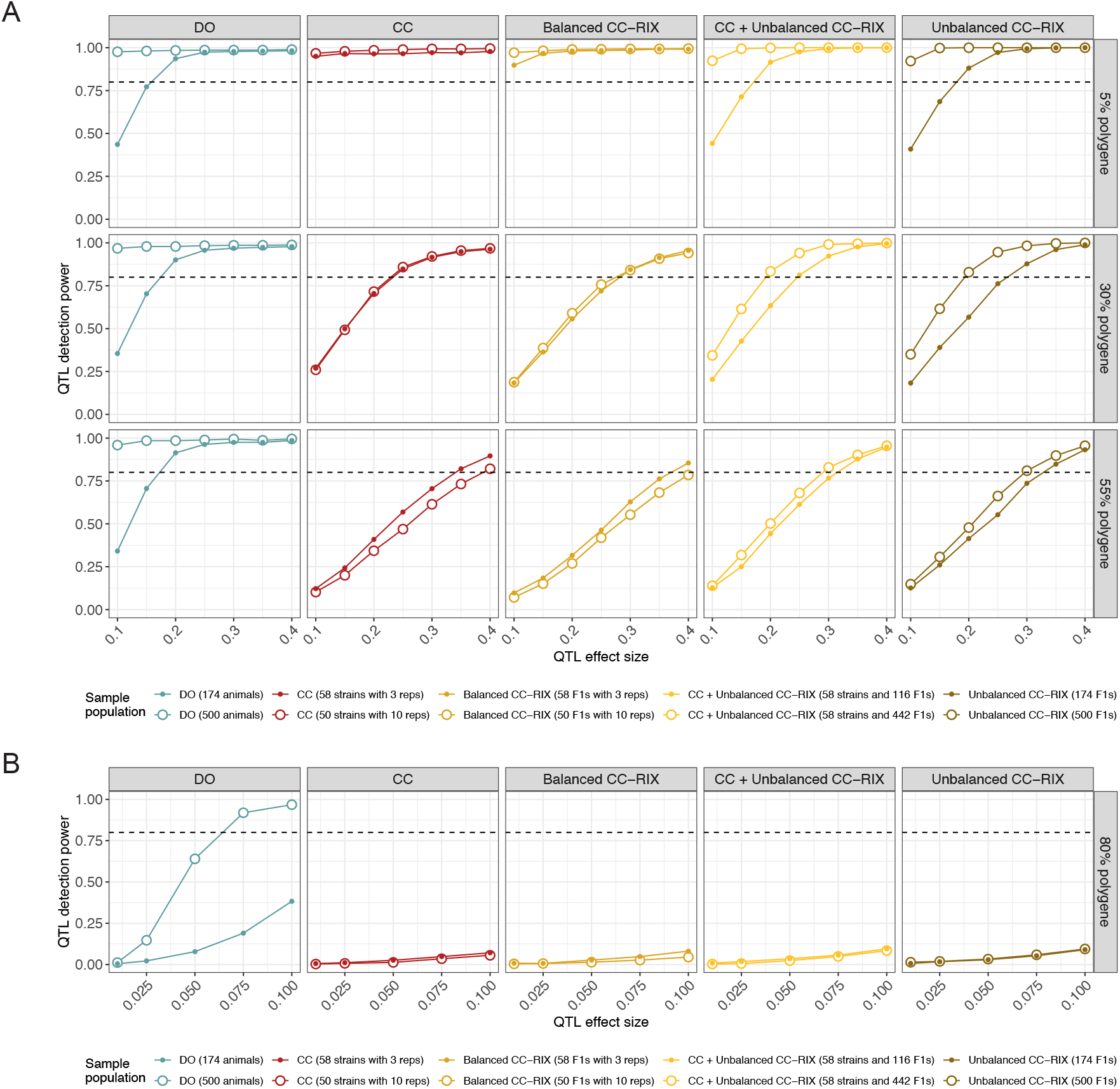
QTL mapping power based on lenient local chromosome significance (such as for detecting *cis*-eQTL) in data simulated for CC, CC-RIX, and DO populations. QTL power curves across populations (columns) and (A) low-to-moderate polygenic background (rows) and (B) a highly polygenic background. Sample populations were composed of 174 (solid points) and 500 mice (open points). Horizontal dashed lines at 80% power included for reference. For power estimates based on stringent genome-wide FWER control, see Figure 6.

### The effect of genotype frequencies on QTL effect size in mapping populations

For the power curves reported above, we strictly scaled the effects of the simulated QTL so that it causes a specified proportion of the total phenotypic variation in the observed mapping population. Consider the case of an allele that is rare in one sample population but common in another; this scaling implicitly increases the effect of the QTL in the population in which the allele is rare to equalize the variation explained across populations. More broadly speaking, controlling the proportion of variation explained by the QTL in the sample population will inflate the QTL effect as its minor allele frequency decreases. In practice this likely inflates the power estimate for the population in which the minor allele is rarer.

In MPPs, allele frequencies will be strongly affected by the allelic series at the QTL, *i*.*e*., how the QTL’s alleles are distributed among the founder strains (Crouse et al. 2020). For all results reported here, we simulated a bi-allelic variant with alleles evenly distributed among the founder strains, which results in fairly balanced allele frequencies across the populations. This assumption is optimistic in terms of power, given that many QTLs are driven by alleles from the phylogenetically distinct CAST and PWK founder strains (Aylor *et al*. 2011; Keele *et al*. 2020). We previously explored how the allelic series of a QTL influenced mapping power in the CC (Keele et al. 2019); less balanced allelic series generally had reduced mapping power due to lower allele frequencies. Researchers should expect QTLs with less balanced allelic series to have reduced power compared to the estimates reported here.

Beyond allele frequencies, when comparing inbred and outbred populations, the genotype frequencies at the QTL can greatly influence the phenotypic variation observed in the sample population. Variation due to the QTL is maximized when the sample population is composed of individuals with homozygous genotypes at the QTL rather than heterozygous, resulting in a larger QTL effect size in the CC than the CC-RIX or DO. Though power curves are often contextualized in terms of the proportion variance explained within the mapping population, given the large-scale differences in genotype frequencies between these populations, we sought to evaluate how these population features would affect mapping power. We again performed 1,000 simulations for QTLs with effect sizes ranging from 10% to 40% (increments of 5%) in a 30% polygenic background for samples of 174 mice. We now scaled the QTL effect equally across the CC, CC-RIX, and DO populations with respect to a reference population with homozygous genotypes and balanced allele frequencies at the QTL. The effect of scaling to the same reference population on mapping power, as well as the relationships between QTL effect size in the sample population and minor allele frequency and heterozygosity are shown in Figure 8.

**Figure 8.**
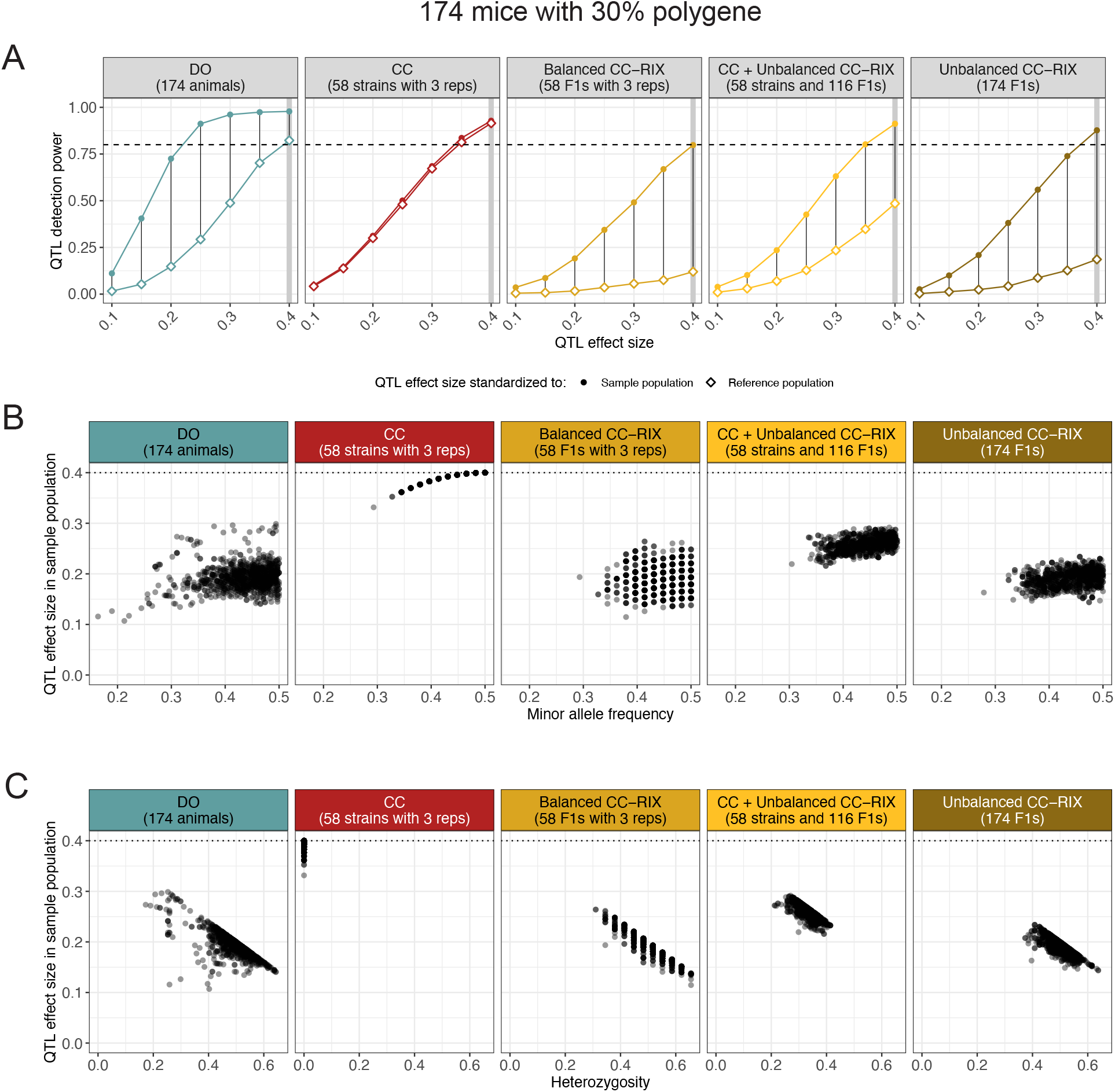
Heterozygosity and allele frequency imbalance reduce the QTL effect size in the sample population and thus QTL mapping power. The data represent 1,000 simulations of QTLs with effect sizes ranging from 10% to 40% in a 30% polygenic background. (A) Reduction in mapping power across all populations when the additive QTL effects are scaled with respect to an inbred population with balanced alleles (reference population, diamond open points) compared to the population being mapped (sample population, circular solid points). Black vertical lines highlight the reduction in mapping power for each simulated QTL effect size. Horizontal dashed lines at 80% power included for reference. Gray vertical bars at 40% QTL effect size are included to highlight simulation setting explored below. (B) Comparison of the observed QTL effect size in the sample population (y-axis) to the QTL minor allele frequency (x-axis) across 1,000 simulated QTLs. Horizontal dashed lines at 40% QTL effect size, representing the QTL effect size in the reference population. (C) Comparison of the observed QTL effect size in the sample population (y-axis) by QTL heterozygosity (proportion of sample population with heterozygous genotype) (x-axis) across 1,000 simulated QTLs. Horizontal dashed lines at 40% QTL effect size, representing the QTL effect size in the reference population.

#### QTL effect size is maximized in the CC

The mapping power in the CC was only slightly reduced based on scaling to the reference population (Figure 8A), due to minor imbalances in allele frequency reducing the QTL effect size (Figure 8B). Heterozygosity had no effect on QTL effect size in the inbred CC because they are fully homozygous (Figure 8C). The DO experienced a larger reduction in mapping power than the CC, resulting in its mapping power being slightly lower than the CC’s, mostly due to large-scale heterozygosity. Mapping powers in the CC-RIX populations were penalized the most, likely due to the combined effects of heterozygosity, imbalanced allele frequencies, and population structure.

These findings suggest there is some cause for optimism when mapping in the CC, in which an additive QTL effect is likely to be maximized within the sample population. It is important to note that deviations from additivity could lessen this effect. These findings also reiterate that the CC-RIX populations have less utility for QTL mapping. Samples of 174 CC and DO are well-powered to detect a large 35% to 40% QTL (in the reference population), whereas none of the CC-RIX have greater than 50% power. We emphasize that though the inbred genotypes of the CC maximize simulated QTL effect size and thus power in some contexts, this benefit will still be limited by the reduced number of unique genomes compared to the DO, particularly as the genetic complexity of the trait increases. Large sample populations of DO are by far the best option for mapping highly polygenic traits.

### QTL location interval performance

QTL location intervals allow researchers to survey and prioritize candidate genetic variants and genes near a QTL. Defining a conventional statistical confidence interval for QTL location is challenging because of the non-smooth likelihood of the location parameter of the QTL model due to discrete genotype markers (Manichaikul *et al*. 2006). Approximations to a confidence interval include likelihood-based methods that estimate intervals using the LOD scores from markers around a detected QTL as a profile likelihood for QTL location. Two such methods include the LOD support (Dupuis and Siegmund 1999) and Bayes credible intervals (Broman and Sen 2009). These approaches were proposed and calibrated in simpler bi-parental crosses (*e*.*g*., F2 intercrosses and backcrosses). The greater complexity of the multi-allelic QTL model (Equation 6) for MPPs compared to bi-parental populations warrants statistical assessment in the context of the CC, CC-RIX, DO.

Sampling the data represents another approach to quantifying uncertainty on QTL location through methods like bootstrapping and permutation. We evaluated three sampling-based QTL location intervals: parametric bootstrap, parametric permutation, and Bayesian bootstrap (Rubin 1981). See Appendix C for more details on these sampling-based methods. Note that in practical terms the likelihood-based approaches are appealing because they require significantly less computation due to no sampling.

We estimated these intervals for the 1,000 simulated QTLs in the CC, CC-RIX, and DO populations of 174 animals in the 40% QTL and 30% polygenic background scenario. For all methods, we evaluated how the QTL coverage rate (the rate that the estimated interval included the true QTL across simulations) compared to the support level or nominal probability. For methods with a nominal probability (*e*.*g*., 95% confidence interval), ideally the observed coverage rate would be close to the nominal probability. We also evaluated methods based on mapping power when strictly requiring the estimated QTL interval to cover the true location.

The support level for a LOD support interval with 80% coverage varied with population and statistical threshold, ranging from 1.25 in the CC and balanced CC-RIX to less than 1 (*≈* 0.7) in the unbalanced CC-RIX populations (Figure S7A). The likelihood-based Bayes credible interval performed similarly to LOD support interval, though power was slightly reduced for all populations (Figure S7B). Bayes credible intervals were also conservative in terms of coverage rate—observed coverage was generally higher than the nominal probability.

#### Likelihood-based QTL intervals outperform sampling-based ones

The sampling-based intervals resulted in reduced mapping power compared to likelihood-based intervals (Figure S8). However, the observed coverage rate for sampling-based methods did generally track more closely with the nominal probability than for the likelihood-based Bayes credible intervals. In terms of coverage rate and the median interval width (narrower being better), the likelihood-based intervals generally performed better (Figures 9 and S9). For likelihood-based methods, the Bayes credible intervals were slightly narrower than the LOD support intervals, but this was balanced by LOD support intervals having slightly better coverage. For sampling-based intervals, parametric bootstrap and permutation outperformed Bayesian bootstrap. In summary, we found likelihood-based intervals to be the superior option for estimating QTL intervals in the CC, CC-RIX, and DO given their computational ease and overall superior statistical performance.

**Figure 9.**
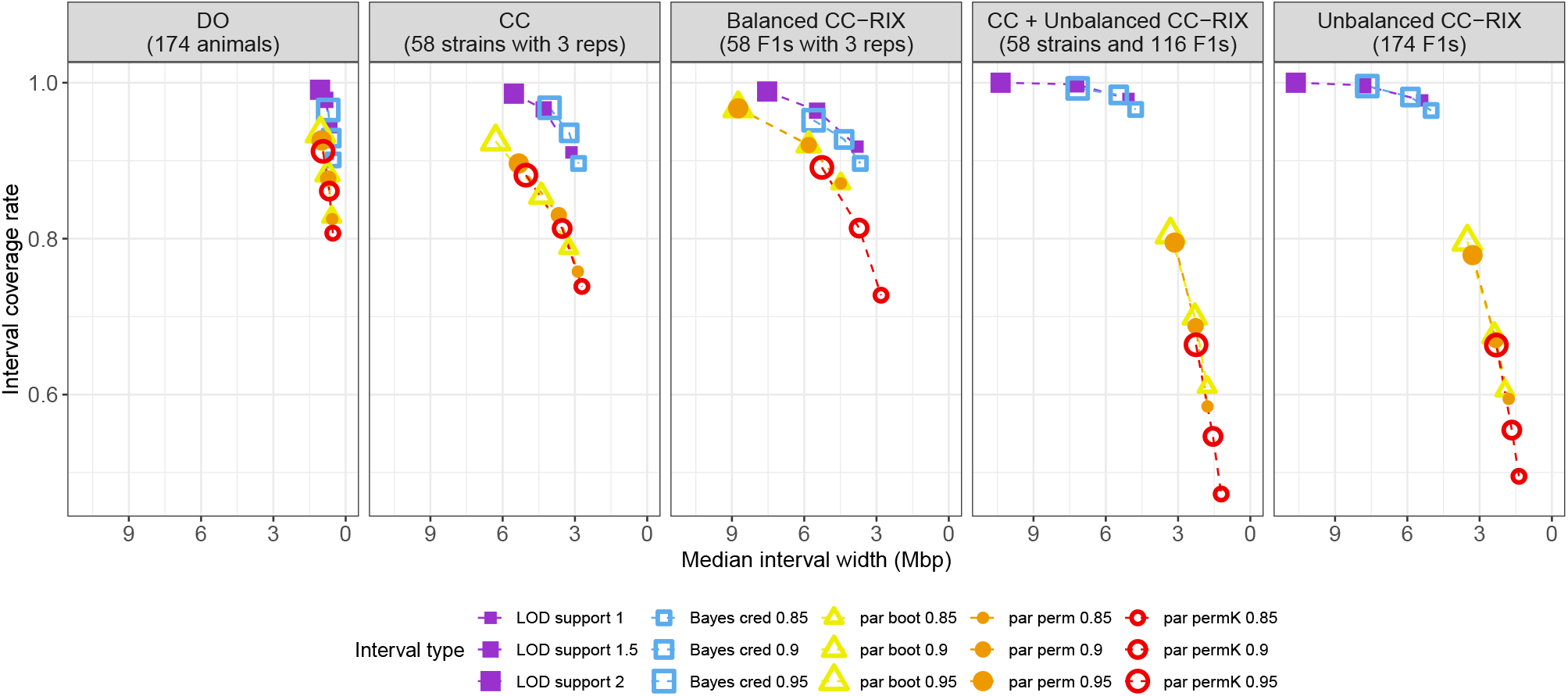
Performance of QTL location intervals across data simulated for CC, CC-RIX, and DO populations in terms of coverage rate (y-axis) and interval width (x-axis). Intervals are summarized over 1,000 QTLs simulated in 174 animals for each sample population (using the 40% QTL and 30% polygenic background setting). Cool colors represent likelihood-based intervals: LOD support (purple) and Bayes credible (Bayes cred; blue). Warm colors represent sampling-based intervals: parametric bootstrap (par boot; yellow), parametric permutation (par perm; orange), and parametric permutation with kinship matrix (par permK; red). For parametric permutation intervals with kinship, 200 samples were generated; for the other sampling-based intervals, 1,000 samples were used. Dashed lines connect summaries from the same procedure but with differing support levels, with increasing support indicated by larger symbols. Summaries for Bayesian bootstrap intervals were excluded due to poor performance, but can be seen in Figure S9.

We note that these results are based on simulated data, which does not often reflect all the structure and complexity of real data. Fitting the founder haplotype model, particularly in smaller sample populations (<200 mice), can result in unstable associations when there are rare alleles present (Keele et al. 2018; Hsiao *et al*. 2020). We speculate that the likelihood-based intervals could be susceptible to these problematic loci, producing overly certain, *i*.*e*., narrow, QTL intervals. In these cases, researchers should consider comparing multiple interval estimates, including sampling-based ones.

### Extending the musppr R package to future studies

We designed musppr to be reusable, allowing researchers to input genetic data from their own sample populations of CC, CC-RIX, and DO, and thus tailoring findings to specific studies. Its functions are amenable to being run in parallel on a computing cluster, allowing deeper evaluations of experimental performance, such as mapping power across more QTL effect size and polygenic background settings, which could be useful for proposals and when planning experiments. The broad findings reported here, such as the value of genetic replicates for estimating heritability, are also largely valid when extrapolating to non-recombinant inbred panels, such as the CC/DO founder strains or the Hybrid Mouse Diversity Panel (Lusis et al. 2016), as well as non-mouse MPPs. We do caveat that to use musppr to analyze genetic data from non-mouse MPPs, functions may need to be expanded or adjusted. For example, if the model underlying the QTL mapping analysis requires population-specific features that are not available in the model fit by the qtl2 R package, musppr’s mapping function would need to be adjusted.

## Conclusions

Here we evaluated the performance of three related genetically diverse mouse MPPs, the CC, CC-RIX, and DO, in estimating heritability and mapping QTLs, commonly used genetic analyses for the study of complex traits. Our findings provide examples of best practices for researchers designing studies with these population resources, such as using the ASV form of the kinship matrix for heritability estimation. More broadly, this work reveals the relative strengths of these populations. Replicate mice in the CC and CC-RIX samples result in more efficient estimation of heritability, potentially offering more precise estimates from far fewer animals than would be required in the DO. The CC and CC-RIX can be powerful tools for genetic mapping when the QTL effect is large (*≥* 40%) and the genetic architecture is fairly simple, but as the trait becomes more polygenic and QTL effect sizes smaller, only large sample populations of DO are likely to be well-powered for QTL mapping. The complex population structure of the CC-RIX reduces mapping power, but does enable more accurate estimation of additive heritability. Furthermore, the CC-RIX can be used to detect parent-of-origin effects using reciprocal F1 designs (Oreper et al. 2018; Sun et al. 2021), though we do not investigate these approaches here. These key principles should extend to MPPs (and more broadly any mapping population) of other organisms with similar experimental features (*e*.*g*., genetic replicates, inbred/outbred).

These results emphasize the complementary nature of these populations for joint analyses. For example, even though CC and CC-RIX sample populations are unlikely to be sufficiently powered to dissect highly polygenic traits, a leniently detected result could confirm a QTL stringently detected in a larger DO sample population; furthermore, replicable CC and CC-RIX mice that possess key alleles of the QTL identified in the DO could be used for follow-up mechanistic studies. Taken together, they represent a flexible and powerful MPP resource for next generation complex trait studies.

## Data availability

All analyses were performed using the R statistical programming language (R Core Team 2022). We wrote the musppr R package to perform all simulations and subsequent analysis, using the qtl2, miQTL, and sommer R packages as dependencies. The musppr R package is available at https://github.com/gkeele/musppr. Founder haplotype data for all populations used in this study, a fixed version of musppr, and R code used to generate the simulated data, figures, and reported results can be found at https://doi.org/10.6084/m9.figshare.20560821.

## Acknowledgments

We acknowledge Martin T. Ferris and William Valdar, both of the University of North Carolina at Chapel Hill, and Gary A. Churchill of the Jackson Laboratory for discussions related to this work. We thank Callan O’Connor of the Jackson Laboratory and Paul L. Maurizio of the University of Chicago for providing feedback on an early draft manuscript.

## Funding

This work was supported by grants from the National Institute of General Medical Sciences (NIGMS) of the National Institutes of Health (NIH): F32GM134599 and R01GM070683.

## Conflicts of interest

The author declares no conflicts of interest.

## Appendix A

### Processing founder haplotype data

Our initial sample populations were 116 CC mice (two animals per 58 strains) genotyped on MiniMUGA (11,000 markers), 192 DO mice genotyped on MegaMUGA (57,000 markers), and 500 DO mice genotyped on GigaMUGA (143,000 markers). We used the qtl2 R package to impute all populations to the same 64,000 loci. Though the shared set of loci far exceeds the mapping resolution of the CC, it is appealing to compare populations based on the same set of loci.

The input data for the DO are scaled haplotype dosages, reflecting an additive model rather than the full 36-state space of outbred diplotypes. Denser genotype arrays result in greater certainty in founder haplotype pair (*i*.*e*., diplotype) estimation. To keep this from influencing our findings, we imputed all loci based on most likely haplotype.

#### CC individuals and strains

Consider diplotype probability vector for CC mouse *i* at the locus *p*:

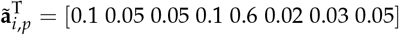

where elements represent the homozygous diplotype probabilities for the founder haplotypes ordered as AJ, B6, 129, NOD, NZO, CAST, PWK, and WSB. For this example, we impute the mouse as having the NZO diplotype:

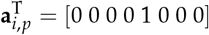

For the CC, we performed this step both at the individual-level and strain-level. For strain-level, we first averaged the diplotype probabilities at all loci between the two individuals from the same strain before doing imputation of most likely diplotype, thus smoothing out loci with segregating variation within a CC strain.

#### CC-RIX

We derived founder haplotype data for the 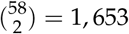 possible F1s given 58 CC strains (ignoring reciprocal F1s). Consider CC strains *j* and *k* that have the B6 and PWK alleles at locus *p*, respectively:

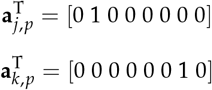

The F1 offspring’s scaled haplotype count at the locus p is calculated by the taking element-wise averages:

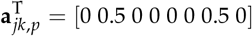

This process was repeated across all loci for all pairwise combinations of CC strains.

#### DO

Selecting most likely haplotypes for the DO samples is more complicated than the CC due to input being scaled haplotype dosages rather than 36-state probibilities. Consider the scaled haplotype dosage vector for DO mouse *i* at the locus *p*, we calculate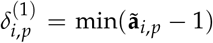 the dosages suggest mouse *i* is homozygous at locus *p* and we set the allele with maximum probability from 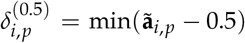 to 1 and all others to 0. If 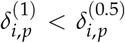, the dosages suggest mouse *i* is heterozygous at locus *p*, and for **a**_*i,p*_ we set the alleles with highest and second highest probability from 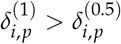 to 0.5 and all others to 0.

## Appendix B

### Simulating data for heritability estimation

We extended our simulation approach for QTL mapping (Keele et al. 2019) to heritability, built from the model specified in Eq 1. We generate simulation *t* using the following model:

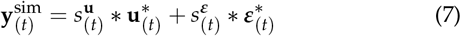

An intercept can be included but by default we set it to 0. We randomly sample draws of 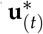 and 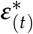 according to 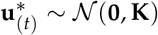 and 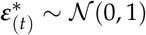. To control the relative contributions of a polygenic effect (**u**) at a specified proportion of total variation *ϕ*^2^, we scale the random draws accordingly:

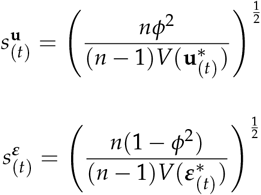

where *V*(.) returns the unbiased sample variance of its argument vector. For simulations with two components of heritability, we used the same approach but with respect to Eq 3, now with two polygenic effects controlled at 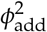 and 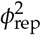.

## Appendix C

### Estimating confidence intervals for the location of a QTL

We estimated likelihood-based intervals for QTL location using LOD support intervals (Dupuis and Siegmund 1999) and Bayes credible intervals (Broman and Sen 2009), which were calculated through the qtl2 R package (Broman et al. 2019). These approaches are fast and efficient because they do not involve a sampling process and are instead summarized directly from the LOD score profile around a detected QTL.

In contrast, sampling-based approaches are slow but statistically appealing because they incorporate uncertainty through a sampling process. We evaluated four related sampling-based procedures.

#### Parametric bootstrap (par boot)

The sampling process for parametric bootstrap involves simulating data from the QTL model (Eq 6) at the detected locus. We fit the model to generate parametric bootstrap sample *t*:

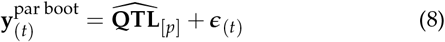

where 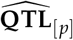 is the fit prediction of the QTL (based on eight fixed effect parameters for founder haplotpes) and ***ϵ***_(*t*)_ is generated according to 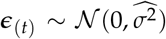 where 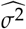is the estimated residual variance. We then map all bootstrap samples for the chromosome of the detected QTL and record the peak location. Middle quantile intervals for peak location were then calculated for a specified level, *e*.*g*., 95%. Notably, we exclude the polygenic effect to simplify the computational cost of running scans of the bootstrap samples. Incorporating variation from estimating the polygenic effect would make the intervals wider.

#### Parametric permutation (par perm)

The sampling process for parametric permutation is similar to parametric bootstrap. To generate parametric permutation sample *t*, we fit:

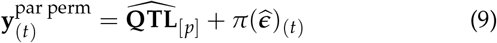

where *π*() is a function that permutes (randomly reorders) a vector, 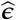 are the fit residuals, and all other terms as previously defined. Subsequent scans of parametric permutations and interval estimation is performed as for the parametric bootstrap.

#### Parametric permutation with kinship (par permK)

We also performed the parametric permutation procedure as previously described but with the kinship term fit as part of the subsequent scans, allowing us to assess whether ignoring the kinship effect makes a significant difference.

#### Bayesian bootstrap (Bayes boot)

The Bayesian bootstrap procedure is a continuous generalization of nonparametric bootstrapping, *i*.*e*., sampling with replacement (Rubin 1981). Instead of sampling with replacement, we generate sample weights for individuals 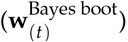, ensuring that all individuals make some contribution to each sample, even if only fractional. The original data are re-scanned for the chromosome with the detected QTL, but now with each sample of weights. Intervals are recorded using the same quantile summary as with the other sampling-based approaches. Initially we included the kinship term but found that when qtl2 was provided a kinship matrix, it disregarded the weights argument. Further work is needed on weighting in the presence of a random term for kinship.

Evaluating the sampling-based procedures represents a significant computational burden because of the additional sampling steps for each simulated QTL. For the 1,000 QTLs that were simulated for the 40% QTL and 30% polygenic background scenario, we evaluated parametric bootstrap and parametric permutation based on 1,000 samples. For parametric permutation with kinship and Bayesian bootstrap, the number of samples was reduced to 200 because both procedures are computationally more intensive.

## Appendix D

### Simulating data for QTL mapping

We simulated QTL data using our previous approach in the CC (Keele *et al*. 2019), now extended for CC-RIX and DO data as well. For sample populations of CC or CC-RIX with replicates, we simulated the data as strain/F1 means, reducing the size of the data and making subsequent mapping analysis more computationally efficient. Consider the QTL model specified in Eq 6 with effects on a trait stemming from the QTL, polygenic background, and random noise. We calculate the expected QTL effect size at the level of strain/F1 means based on the reduction in noise on means. Similar to our simulations of heritability (Appendix B), we generate simulation *t* for a QTL at *p* with the following model:

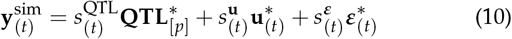

where the *∗* superscript indicates randomly sampled vectors, 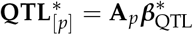, and scaling factors are calculated as

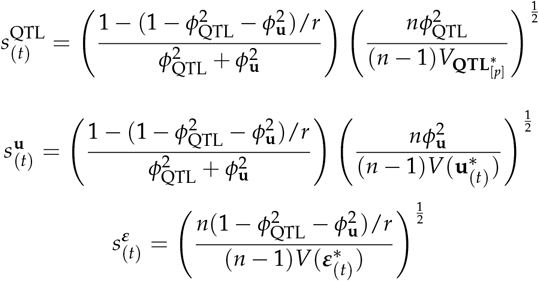

for the proportions of variation explained by the QTL.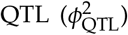 and polygenic background 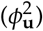, *r* is the number of replicates per strain/F1, 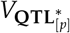 is a variance for the QTL term, and all other terms as previously defined.

We evaluated QTL mapping power based on scaling the QTL effect with respect to both the sample population and a shared reference population that has homozygous and balanced allele frequencies at the QTL. Which scaling is used is determined by adjusting the 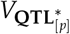 term. When scaling with respect to the sample population, 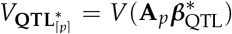. If scaling with respect to the reference population, then 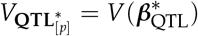.

## Supplemental Figures

**Figure S1.**
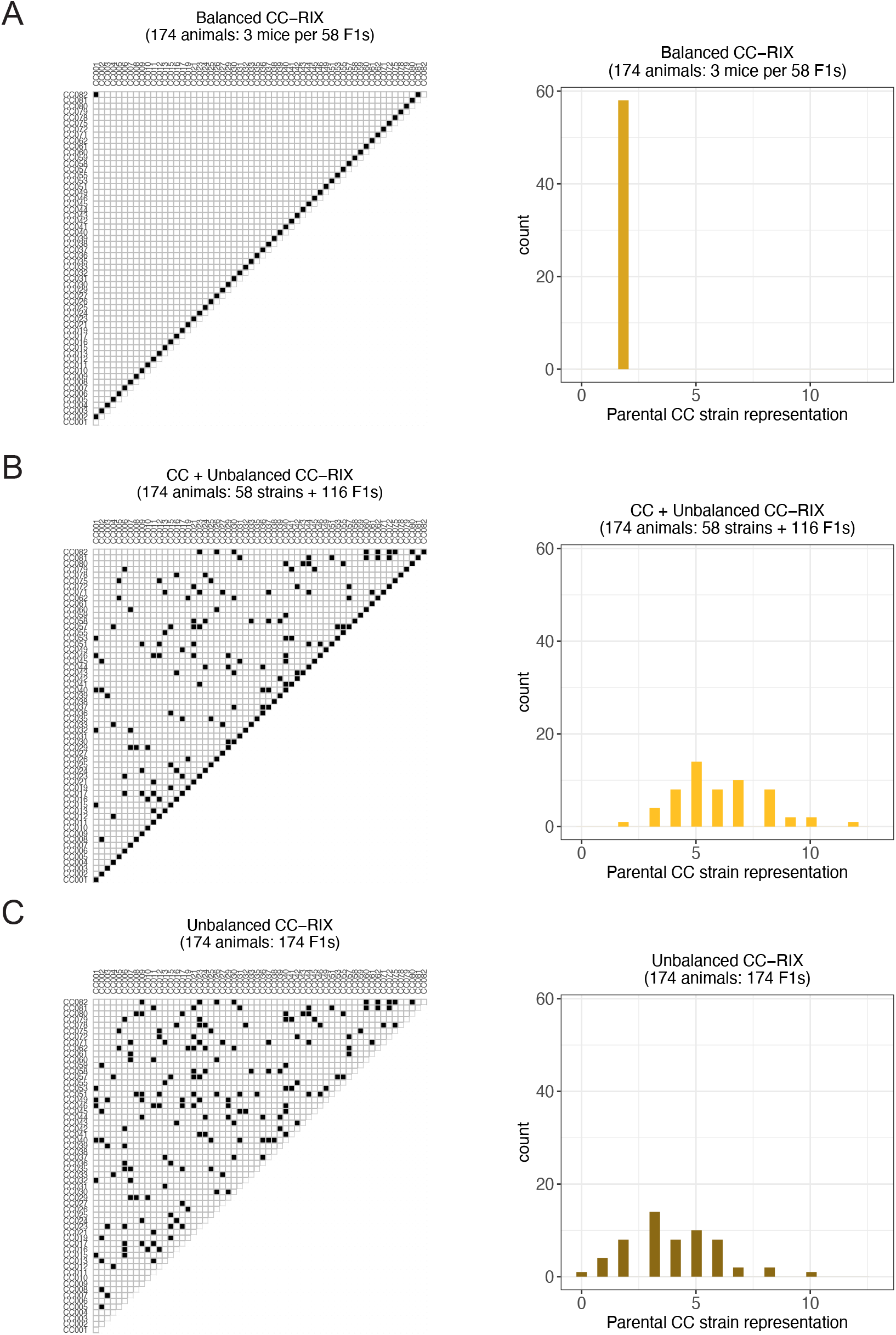
Examples of CC-RIX sample populations (as used in Figure 3C-E): (A) a balanced F1 set where each CC parental strain is equally represented twice, (B) an unbalanced F1 set combined with the CC strains, and (C) an unbalanced set of CC-RIX F1s alone. Each population is represented with a diallel grid of the CC strains and their F1s (left column) and a histogram of the how many F1s/strains represent each parental CC strain (right column). Selected strains and F1s are denoted as black cells in the diallel grid. The effects of sex chromosomes and parent-of-origin are ignored.

**Figure S2.**
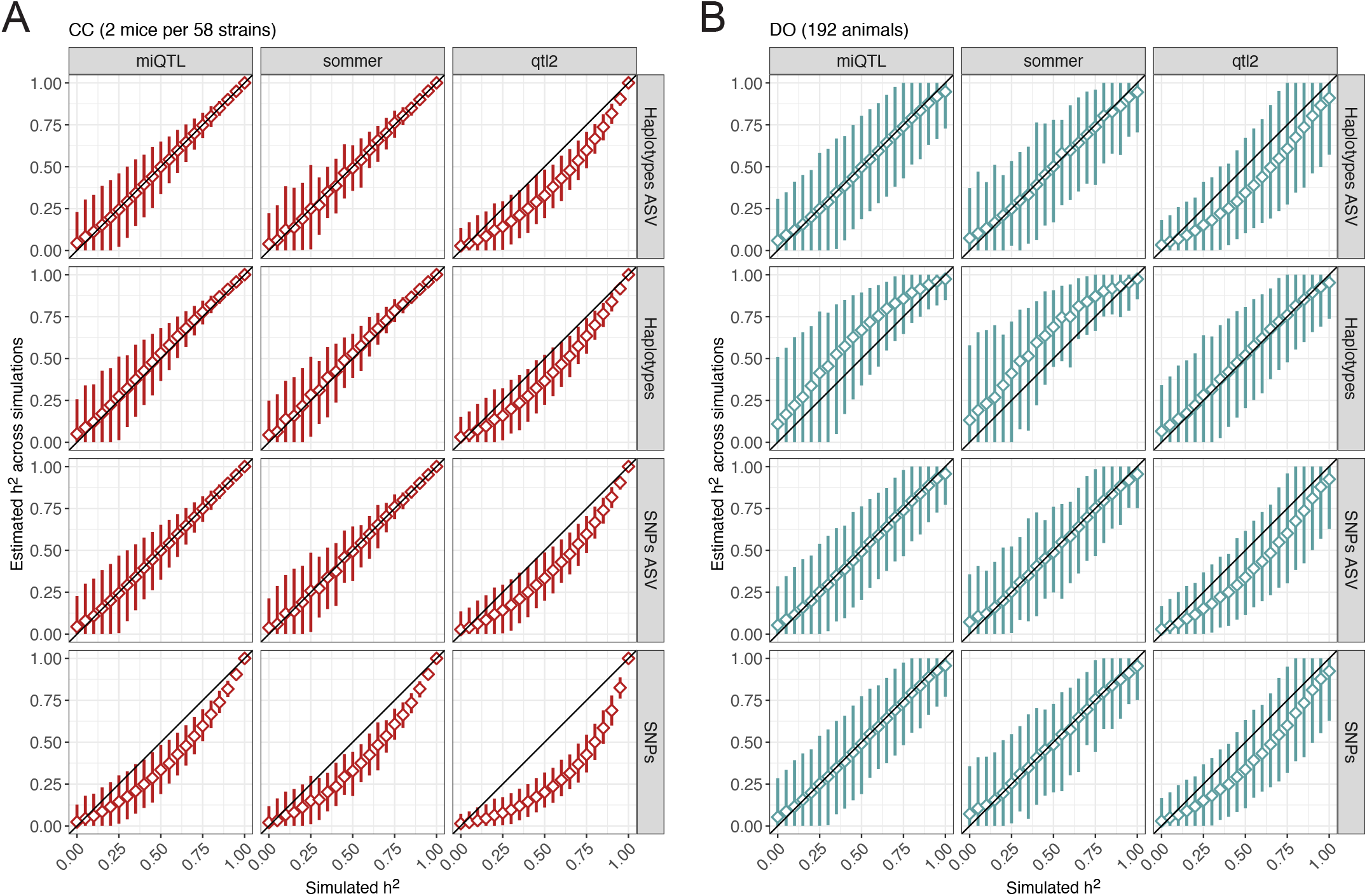
Performance of heritability estimation across kinship matrix estimates (rows) and software packages (columns) in data simulated for (A) 116 CC mice (two per strain) and (B) 192 DO mice. Diamonds represent the mean estimated heritability across simulations from the true heritability. 1,000 simulations were performed with qtl2 and miQTL and 100 were performed for sommer. Vertical line segments represent middle 95% intervals across the simulations. Black diagonal lines indicating the heritability estimate is equal to the true value (y = x) included for reference.

**Figure S3.**
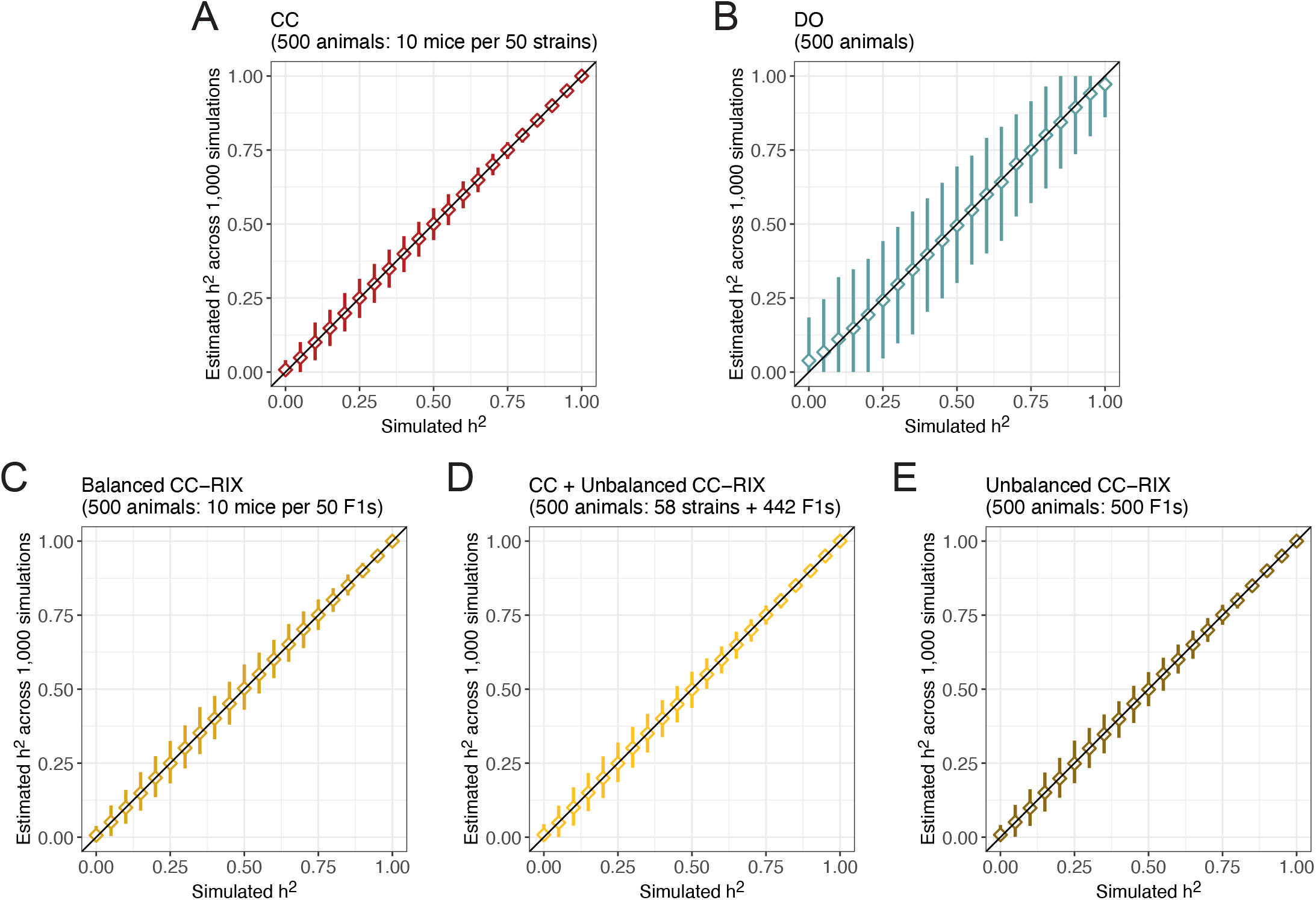
Performance of heritability estimation in data simulated for 500 mice from (A) CC, (B) DO, and (C-E) CC-RIX populations. Diamonds represent the mean estimated heritability across 1,000 simulations from the true heritability. Vertical line segments represent middle 95% intervals across the 1,000 simulations. Black diagonal lines indicating the heritability estimate is equal to the true value (y = x) included for reference. See Figure 3 for results from simulations of 174 mice.

**Figure S4.**
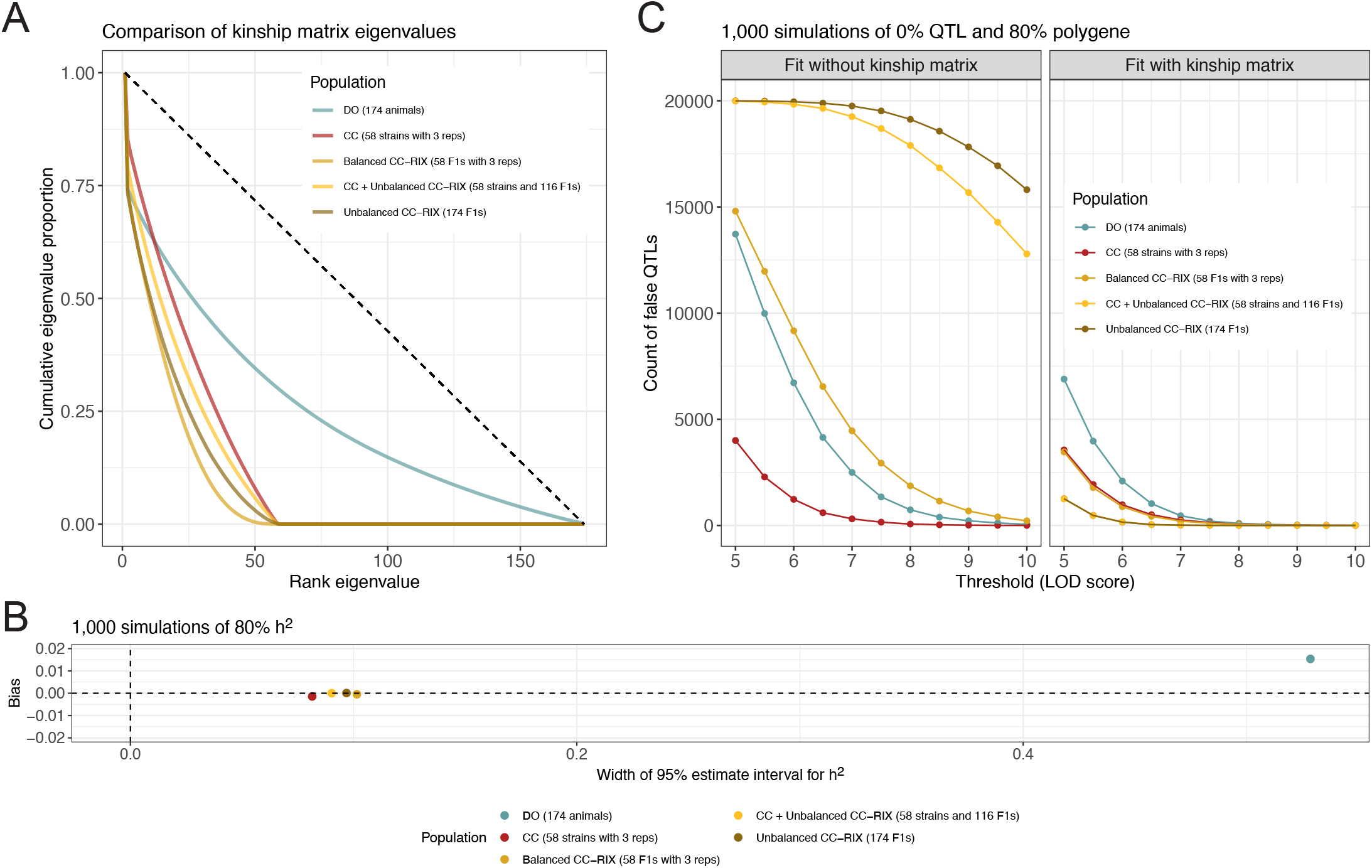
Population structure or unequal relatedness influences heritability estimation and QTL mapping. (A) Population structure summarized as cumulative eigenvalues (y-axis) by the rank of the eigenvalues of the kinship matrices for the CC, CC-RIX, and DO populations. Diagonal dashed line included for reference representing a population with no structure. (B) Comparison of heritability estimation bias (y-axis) to its precision in the form of 95% estimate interval width (x-axis) across the populations. Summaries for each population are based on 1,000 simulations with heritability set to 80%. Horizontal and vertical dashed lines at 0 included for reference representing a perfect summary with no bias or uncertainty. (C) Population structure can result in false positive QTLs when it is not accounted for in the model. The number of false QTLs were counted from 1,000 simulations of data with no QTL and 80% polygenic effect size for all populations.

**Figure S5.**
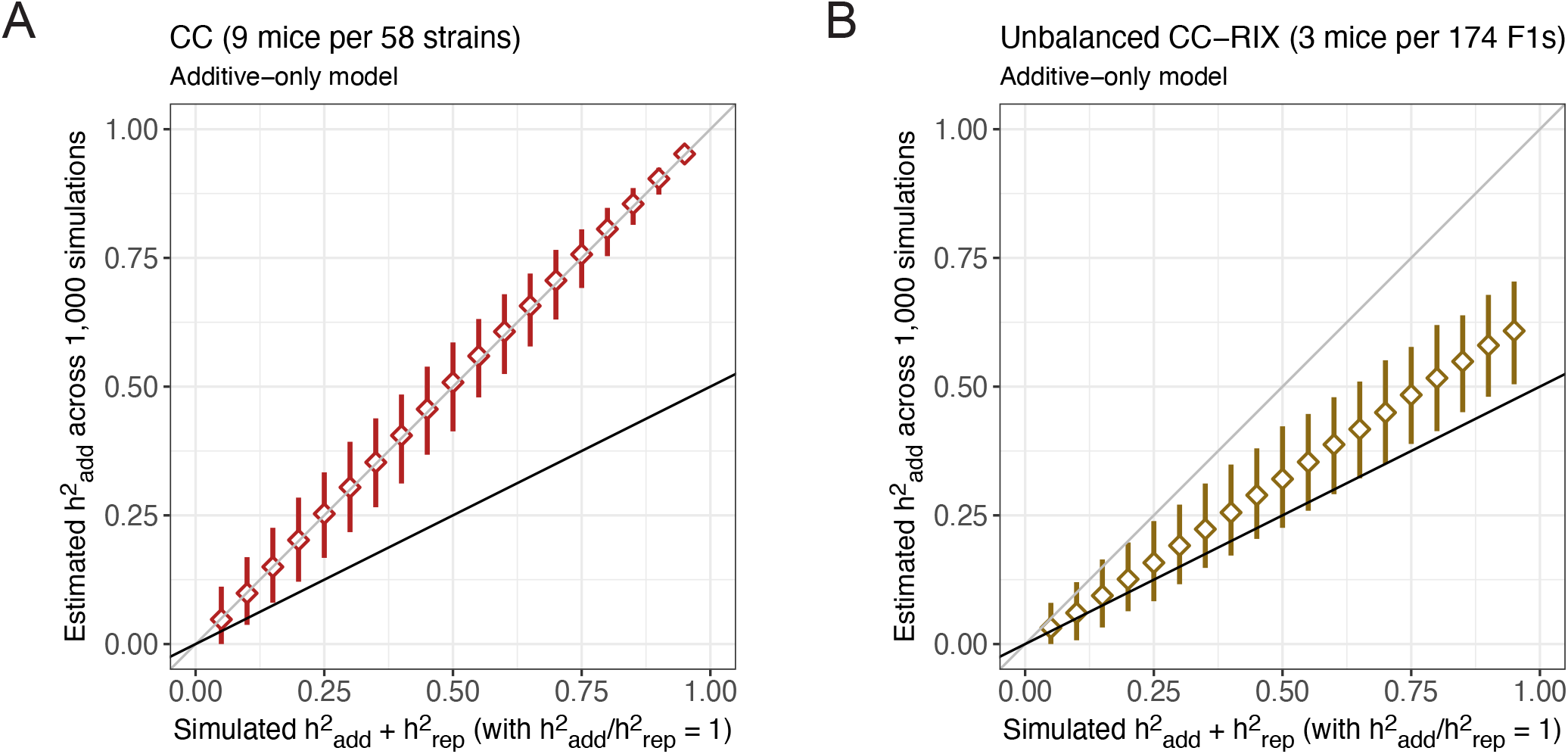
Additive and strain-specific genetic effects are confounded in the CC, but less so in the CC-RIX. Performance of heritability estimation in data simulated for 522 mice from (A) CC and (B) CC-RIX populations with additive 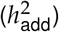 and strain/F1 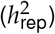 components but the fit model is miss-specified with only an additive component. Diamonds represent the mean estimated heritability across 1,000 simulations from the true heritability. Vertical line segments represent middle 95% intervals across the 1,000 simulations. Black lines indicating the additive heritability component estimate is equal to the true value (y = 2x) included for reference. Gray lines indicating the additive heritability component estimate is equal to the true value of the sum of heritability components (y = x) included for reference.

**Figure S6.**
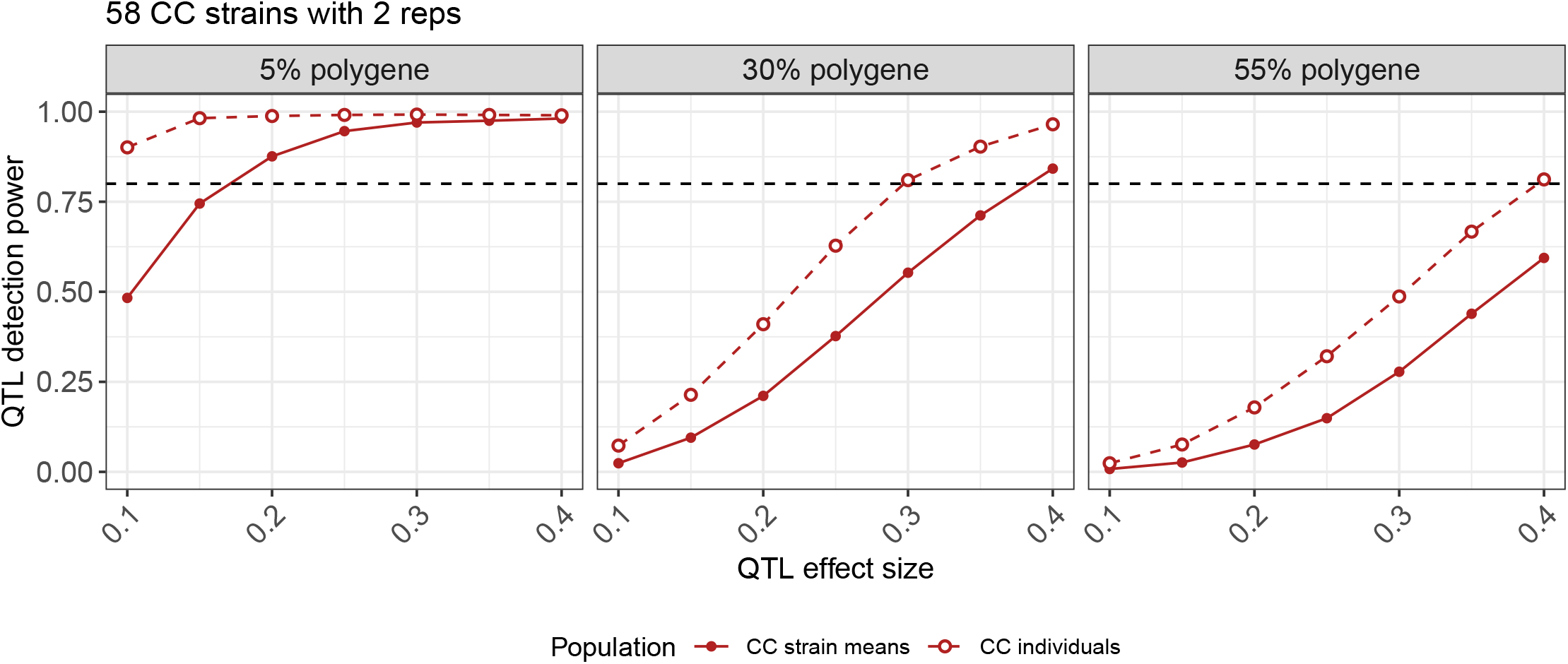
Comparison of QTL mapping power between mapping on CC strain means or the individual-level data. Power was summarized over 1,000 simulated QTLs using genome-wide significance thresholds across low-to-moderate polygenic backgrounds (columns). Simulations were of 116 animals (two animals per 58 CC strains). Horizontal dashed lines at 80% power included for reference.

**Figure S7.**
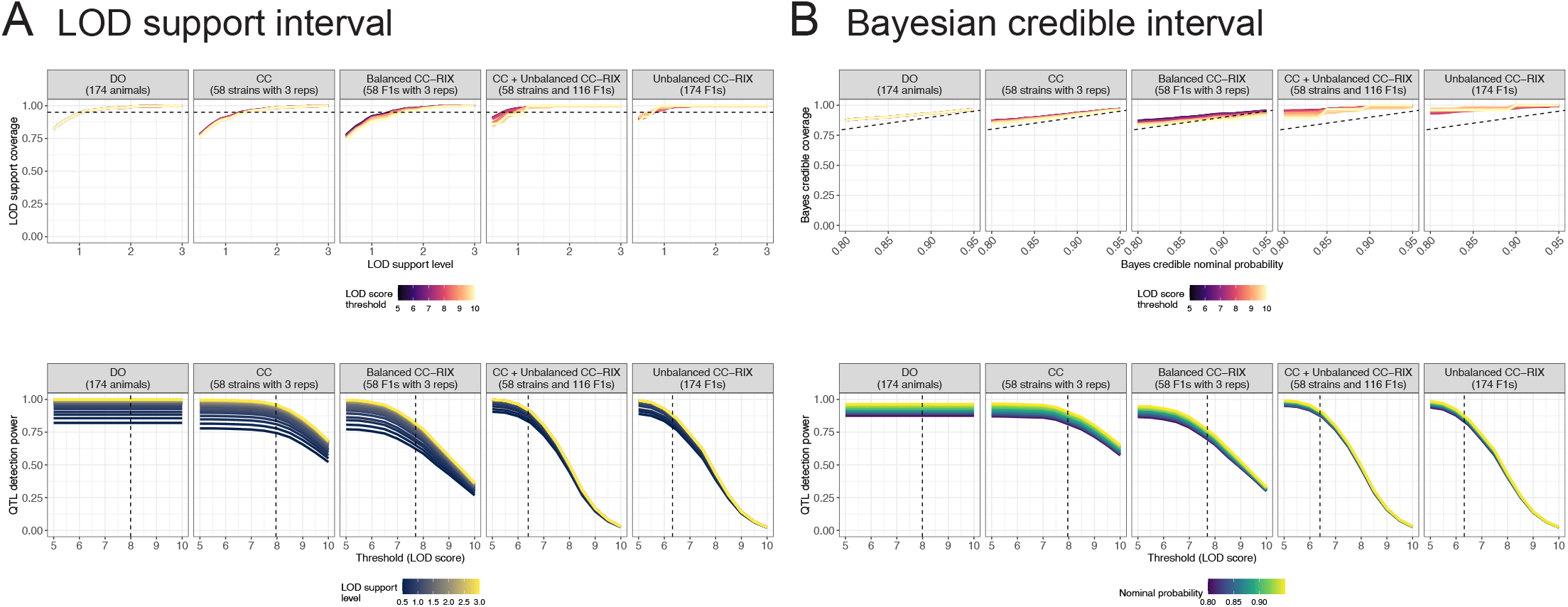
Performance of likelihood-based intervals for QTL location for CC, CC-RIX, and DO populations in terms of (top) QTL coverage rate and (bottom) mapping power. Likelihood-based approaches included (A) LOD support intervals and (B) Bayes credible intervals. Intervals are summarized over 1,000 QTLs (40% QTL with 30% polygenic background) simulated in 174 animals for each sample population. For the coverage rate of LOD support intervals, horizontal dashed lines at 80% coverage included for reference. For the coverage rate of Bayes credible intervals, diagonal dashed lines indicating the interval coverage rate is equal to the nominal probability (y = x) included for reference. For mapping power, vertical dashed lines represent the genome-wide 95% significance thresholds for each population. See Figure S8 for results from sampling-based intervals.

**Figure S8.**
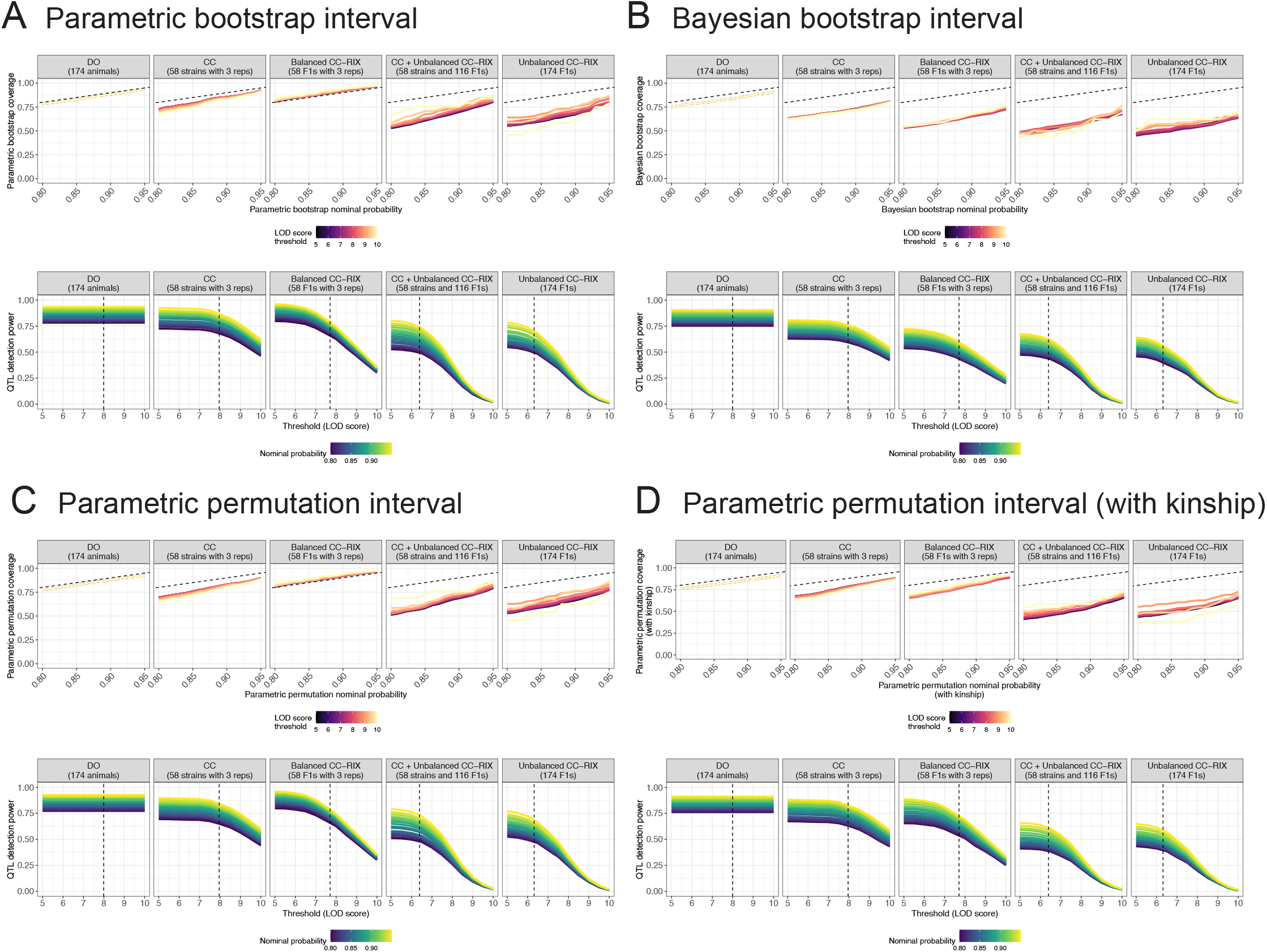
Performance of sampling-based intervals for QTL location for CC, CC-RIX, and DO populations in terms of (top) QTL coverage rate and (bottom) mapping power. Sampling-based approaches included (A) parametric bootstrap intervals, (B) Bayesian bootstrap intervals, (C) parametric permutation intervals, and (D) parametric permutation intervals with kinship included. Intervals are summarized over 1,000 QTLs (40% QTL with 30% polygenic background) simulated in 174 animals for each sample population. For Bayesian bootstrap and parametric permutation intervals with kinship, 200 samples were generated; for the other interval types, 1,000 samples were used. For coverage rate, diagonal dashed lines indicating the interval coverage rate is equal to the nominal probability (y = x) included for reference. For mapping power, vertical dashed lines represent the 95% significance thresholds for each population. See Figure S7 for results from likelihood-based intervals.

**Figure S9.**
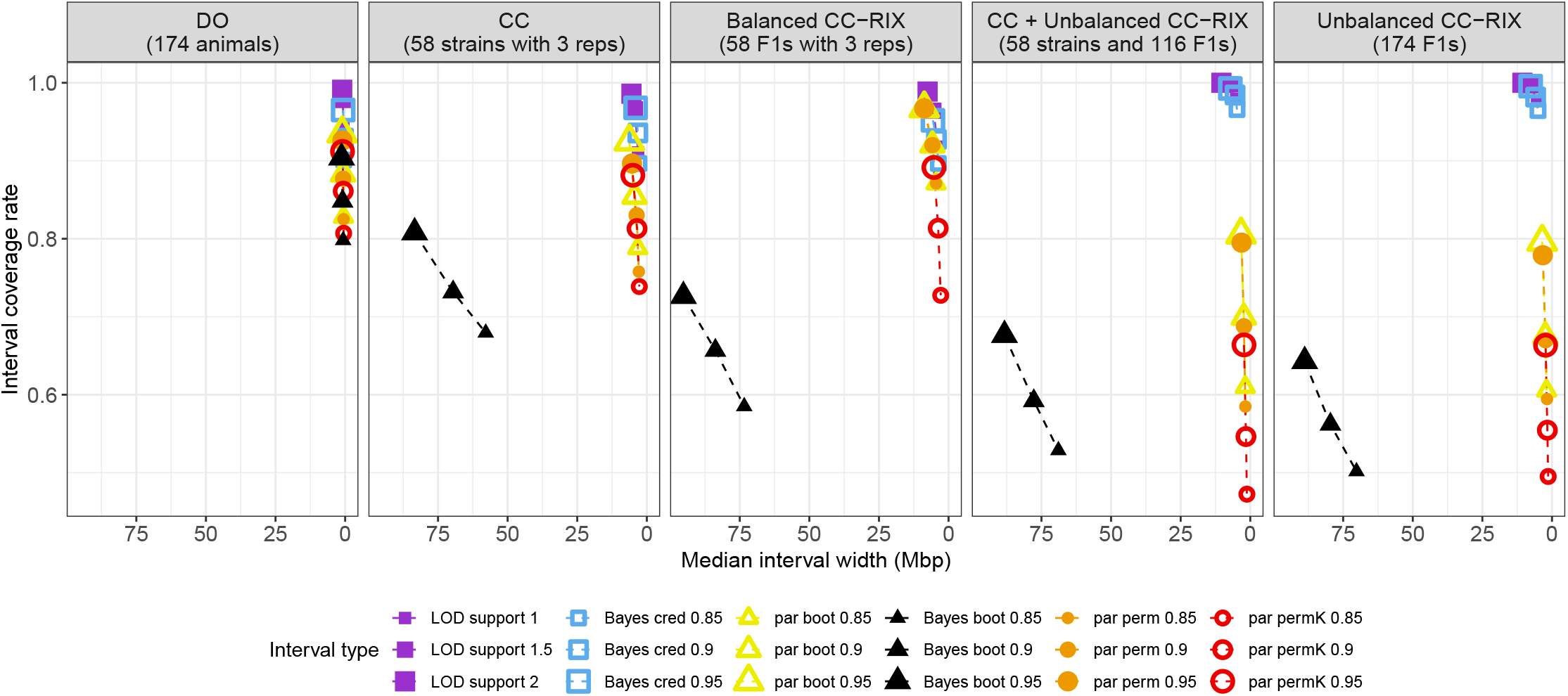
Performance of QTL location intervals across data simulated for CC, CC-RIX, and DO populations in terms of QTL coverage rate (y-axis) and interval width (x-axis). Intervals are summarized over 1,000 QTLs simulated in 174 animals for each sample population (using the 40% QTL and 30% polygenic background setting). Cool colors represent likelihood-based intervals: LOD support (purple) and Bayes credible (Bayes cred; blue). Warm colors represent sampling-based intervals: parametric bootstrap (par boot; yellow), parametric permutation (par perm; orange), and parametric permutation with kinship matrix (par permK; red). Another sampling-based interval, Bayesian bootstrap (Bayes boot) is colored black. Dashed lines connect summaries from the same procedure but with differing support levels, with increasing support indicated by larger symbols. For Bayesian bootstrap intervals and parametric permutation intervals with kinship, 200 samples were generated; for the other sampling-based intervals, 1,000 samples were used. The same summaries are shown in Figure 9 with Bayesian bootstrap intervals omitted to increase clarity.

## Notes

https://doi.org/10.6084/m9.figshare.20560821

### Competing Interest Statement

The authors have declared no competing interest.

### Summary of Updates

The QTL mapping simulations have been greatly expanded to produce power curves across various settings. Figure 4 is revised. A new Figure 7 and Figure S1 were added. Overall text has been updated to improve clarity.

